# Mouse Pachytene piRNAs Cleave Hundreds of Transcripts, But Alter the Steady-State Abundance of Only a Minority of Targets

**DOI:** 10.1101/2024.11.02.621675

**Authors:** Katharine Cecchini, Nandagopal Ajaykumar, Ayca Bagci, Phillip D. Zamore, Ildar Gainetdinov

## Abstract

In animals, 18–35-nt piRNAs guide PIWI proteins to regulate complementary RNAs. During male meiosis, mammals produce an exceptionally abundant class of piRNAs called pachytene piRNAs. Pachytene piRNAs are required for spermatogenesis and have been proposed to control gene expression by various mechanisms. Here, we show that pachytene piRNAs regulate targets predominantly, if not exclusively, by endonucleolytic cleavage. Remarkably, pachytene piRNAs slice hundreds of RNAs, yet a change in steady-state level is detectable for a small fraction of transcripts. Our data suggest that cleavage of the few targets whose abundance is reduced significantly by piRNAs is essential for male fertility. Other pachytene piRNA targets are enriched for highly transcribed genes, which may explain why piRNA cleavage is often inconsequential for the steady-state abundance of targets. We propose that the retention of pachytene piRNAs throughout mammalian evolution is driven by the selective advantage conferred by a tiny minority of piRNAs.

## INTRODUCTION

PIWI-interacting RNAs (piRNAs) are an animal-specific class of small RNAs that silence transposons, regulate host genes, and repress viral transcripts.^1–4^ piRNAs direct PIWI proteins to cleave complementary RNAs in the cytoplasm or recruit transcriptional repressors to nascent RNAs in the nucleus. Most animals produce piRNAs both in germ cells and in somatic tissues.^5–7^ Some arthropods and all vertebrates appear to have lost somatic piRNAs and only make piRNAs in the germline.^5,8^

Unlike microRNAs (miRNAs) and small interfering RNAs (siRNAs), which derive from double-stranded RNAs, piRNAs are produced from long single-stranded precursors.^9,10^ piRNA precursors are transcribed from dedicated genomic loci called piRNA clusters.^11–16^ For example, in fly ovaries and mouse fetal testes, transposon-targeting piRNAs are made from piRNA precursors rich in transposon sequences.^13,17^

At the onset of male meiosis, placental mammals produce pachytene piRNAs, a distinct piRNA class made from a subset of testis-specific long non-coding RNAs (lncRNAs) devoid of active transposon sequences.^11,12,14,16,18^ Pachytene piRNAs are extraordinarily abundant (∼10 million piRNAs / primary spermatocyte) and their biogenesis involves a feedback amplification mechanism: pachytene piRNA-directed cleavage of precursor transcripts initiates the production of more pachytene piRNAs.^19–21^ Of the 100 mouse pachytene piRNA source loci, just four have been genetically disrupted, and only two of these are required for normal male fertility.^20,22–24^

The loci producing the majority of pachytene piRNAs are present at syntenic locations in all placental mammals, yet their primary sequences are not conserved.^16^ In fact, the sequences of pachytene piRNA-producing loci diverge among species nearly as rapidly as non-transcribed regions of the genome.^16^

The majority of pachytene piRNAs are extensively complementary only to the genomic loci from which they are transcribed. The targets of pachytene piRNAs are thus not obvious, and several models have been proposed to explain their function. Pachytene piRNAs have been reported to (1) destabilize partially complementary transcripts by a miRNA-like mechanism^25^; (2) cleave extensively complementary RNAs by an siRNA-like mechanism^20,24,26,27^; (3) activate translation of partially complementary mRNAs^28,29^; and (4) lack intrinsic function and instead be the degradation products of lncRNAs.^30^

Here, we report that mouse pachytene piRNAs regulate targets exclusively by endonucleolytic cleavage. Only several hundred pachytene piRNAs (∼1%) have at least one transcript with sufficient complementarity to direct its slicing. Remarkably, cleavage by most pachytene piRNAs does not change the steady-state levels of their targets. Our analyses suggest that production of fully functional sperm requires cleavage of the few targets whose abundance is changed by piRNAs. We find that other pachytene piRNA targets are typically from highly transcribed genes, potentially explaining why piRNA-directed cleavage often has little impact on target steady-state abundance. Because pachytene piRNAs that change the steady-state levels of targets are rare, we propose that such piRNAs are more often deleterious than advantageous and are removed by purifying selection. Our data suggest that the fitness advantage provided by a minority of pachytene piRNAs ensures that the non-functional majority of pachytene piRNAs are retained in mammalian evolution.

## RESULTS

### Individual Pachytene piRNA Loci are Required for Production of Fully Functional Sperm

Genetic disruption in mice of any of the four largest sources of pachytene piRNAs, *2- qE1-35981(+)* (“*pi2*”), *7-qD2-24830(−)11976(+)* (“*pi7*”), *9-qC-31469(−)10667(+)* (“*pi9*”), or *17-qA3.3-27363(−)26735(+)* (“*pi17*”) has no detectable effect on male fertility or sperm count (Figures 1A, 1B, and S1A; refs.^20,22,23^). Nonetheless, *pi2^−/−^*, *pi9**^−/−^***and *pi17**^−/−^*** single mutants displayed impaired sperm motility (Figures 1C, S1B, and S1C). *pi2^−/−^*, *pi9^−/−^ and pi17^−/−^*sperm showed defects in hyperactivation, the hallmark of Ca^2+^- induced sperm capacitation: the fraction of hyperactivated sperm was halved in *pi2^−/−^*, *pi9^−/−^* and *pi17^−/−^* males compared to C57BL/6 controls (C57BL/6, median 9.2%; *pi2^−/−^*, median = 5.8%, unpaired, two-tailed Mann-Whitney test, Benjamini-Hochberg [BH] corrected *p*-value = 0.008 vs. control; *pi9^−/−^*, median = 4.7%, *p* = 0.003; *pi17^−/−^*, median = 4.5%, *p* = 0.008; Figure 1C). Moreover, *pi2^—/—^* and *pi9^—/—^* single mutants had fewer progressively motile sperm (C57BL/6, median = 6.9%; *pi2^−/−^*, median = 4.4%, *p* = 0.01; *pi9^−/−^*, median = 3.8%, *p* = 0.0005; Figure S1C). These data demonstrate the importance of individual pachytene piRNA loci in the production of fully functional sperm.

**Figure 1.**
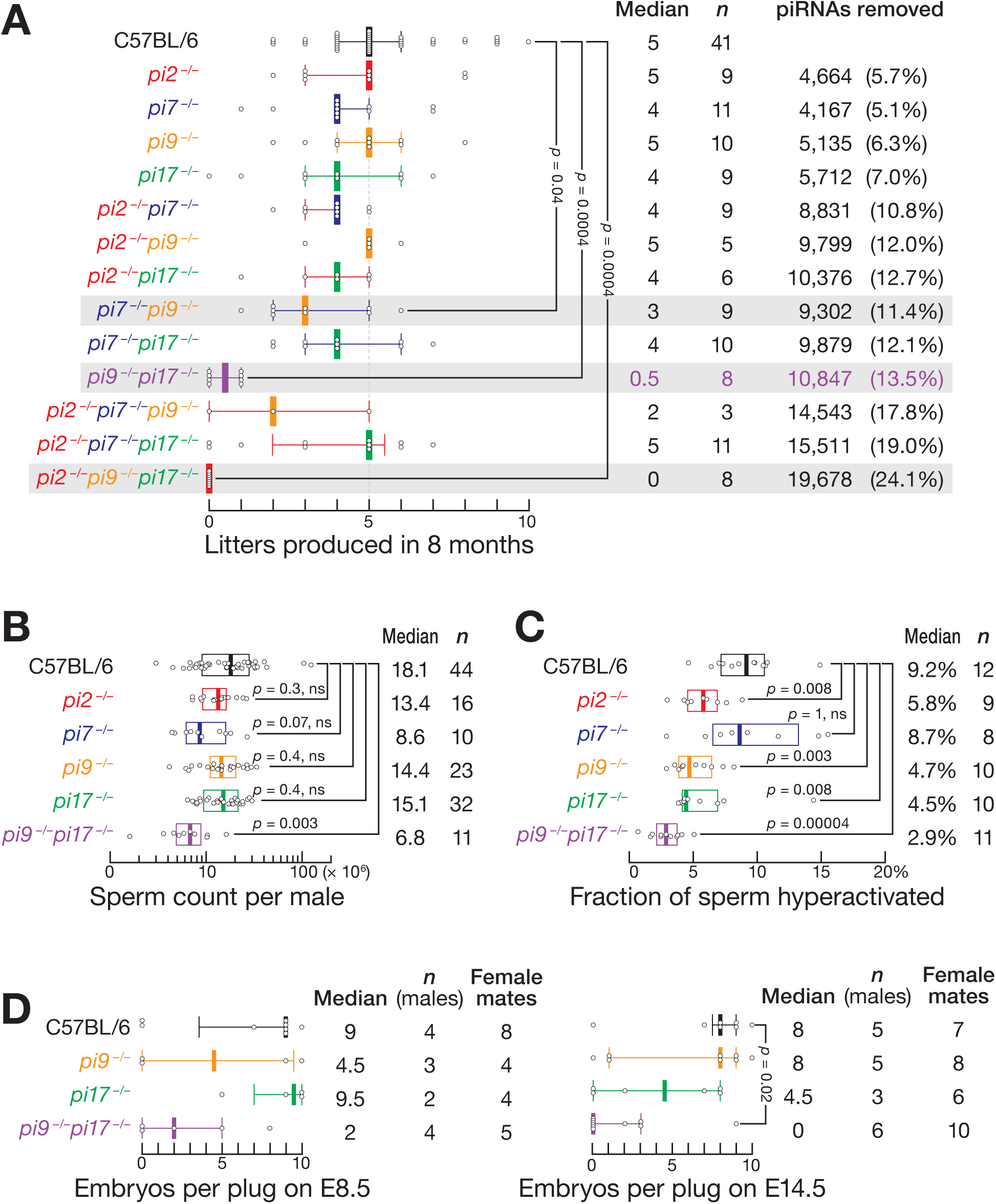
Male Mice Lacking Two or Three Pachytene piRNA Loci Display Fertility Defects. (A) Number of litters produced by the control (C57BL/6) and pachytene piRNA mutant males in successive matings with C57BL/6 females over 8 months. Median and inter-quartile range (IQR) are shown. Kruskal-Wallis test (one-way ANOVA on ranks) *p*-value = 7.3×10^−7^. Benjamini-Hochberg corrected *p*-values for *post hoc* pairwise Mann-Whitney tests are shown. (B) Total number of sperm in caudal epididymis of C57BL/6, *pi9^−/−^*, *pi17^−/−^*, and *pi9^−/−^pi17^−/−^* males. Kruskal-Wallis test (one-way ANOVA on ranks) *p*-value = 0.00087. Benjamini-Hochberg corrected *p*-values for *post hoc* pairwise Mann-Whitney tests are shown. (C) Fraction of hyperactivated sperm from caudal epididymis of C57BL/6, *pi9^−/−^*, *pi17^−/−^*, and *pi9^−/−^pi17^−/−^* males determined by CASAnova. Kruskal-Wallis test (one-way ANOVA on ranks) *p*-value = 7.9×10^−6^. Benjamini-Hochberg corrected *p*-values for *post hoc* pairwise Mann-Whitney tests are shown. (D) Number of embryos produced by males mated with C57BL/6 females at embryonic day (E) 8.5 or E14.5 after mating. For E14.5, Kruskal-Wallis test (one-way ANOVA on ranks) *p*-value = 0.044l; Benjamini-Hochberg corrected *p*-values for *post hoc* pairwise Mann-Whitney tests is shown. For E8.5, non-parametric Kruskal-Wallis and Mann-Whitney tests do not detect difference due to lower number of replicates.

To determine whether the absence of a fertility phenotype for individual mutants reflects genetic redundancy, we generated all possible double and triple homozygous mutant males and tested their fertility in successive matings with C57BL/6 females. Remarkably, one double-mutant combination, *pi9^−/−^pi17^−/−^* caused male infertility (Figures 1A and S1A). A second double-mutant, *pi7^−/−^pi9^−/−^*, showed reduced fertility: the *pi7^−/−^pi9^−/−^* males sired a median of three litters during eight months, compared to five for control C57BL/6 males (Figures 1A and S1A). The other four double mutants were either fully fertile or weakly subfertile. The triple-mutant *pi2^−/−^pi9^−/−^pi17^−/−^*was sterile and the *pi2^−/−^pi7^−/−^pi9^−/−^* was subfertile, as expected (Figures 1A and S1A). Yet *pi2^−/−^pi7^−/−^pi17^−/−^*, which lacks 19% of all pachytene piRNAs, corresponding to >15,500 different piRNA sequences, was as fertile as C57BL/6 control. Together, these data suggest extensive genetic redundancy among the major pachytene piRNA-producing loci in male mice.

### *pi9^−/−^pi17^−/−^* Sperm Displays Defects in Motility, Capacitation, and Fertilization

*pi9^−/−^pi17^−/−^* males were infertile (Figures 1A, 1D, and S1A). The median number of embryos carried by C57BL/6 females at day 14.5 after mating with *pi9^−/−^pi17^−/−^*males was 0, compared to 8 embryos for C57BL/6 females that mated with C57BL/6 males (unpaired, two-tailed Mann-Whitney test, BH-corrected *p*-value = 0.02; Figure 1D). In fact, in 8 months, *pi9^−/−^pi17^−/−^* males sired only 0–1 litters versus 2–10 litters for the C57BL/6 controls (*p*-value = 0.0004; Figure 1A). Consistent with their reduced fertility, *pi9^−/−^pi17^−/−^*produced ∼3-fold fewer sperm than C57BL/6 males (unpaired, two-tailed Mann-Whitney test, BH-corrected *p*-value = 0.003; Figure 1B) and had fewer motile, progressive, and hyperactivated sperm (Figures 1C, S1B, and S1C).

*pi9^−/−^pi17^−/−^* sperm were inefficient in penetrating the outer oocyte layer, the zona pellucida (Figure S1D); the same defect was previously observed for sperm from males lacking piRNAs from a locus on chromosome 6, *6-qF3–28913(−),8009(+)* (“pi6”; ref.^20^). With zona pellucida intact, only 10 ± 10% of oocytes incubated with *pi9^−/−^pi17^−/−^*sperm reached the two-cell stage (703 total oocytes; *n* = 3), compared to 70 ± 30% for C57BL/6 (783 total oocytes; *n* = 3; Figure S1D). However, in vitro fertilization rates were indistinguishable when zona pellucida was removed: 80 ± 30% for *pi9^−/−^pi17^−/−^*(438 total oocytes; *n* = 3) and 80 ± 40% for C57BL/6 sperm (468 total oocytes; *n* = 3; Figure S1D). We conclude that *pi9* and *pi17* piRNAs are required for successful spermatogenesis and oocyte fertilization.

### *pi9* and *pi17* piRNAs Regulate Genes that Control the DNA Damage Response, Apoptosis, and Cell Proliferation

The *pi9^−/^*^−^ and *pi17^−/−^*mutants were generated using a pair of sgRNAs to guide Cas9-mediated deletion of promoter sequences. To exclude Cas9-induced off-target changes in gene expression, we used two different pairs of sgRNAs to generate two independent alleles for each locus (Table S1). In this study, we consider only molecular changes detected in both alleles and therefore refer to the mutations as *pi9^−/−^ and pi17^−/−^*.

Although *pi9^−/−^* and *pi17^−/−^* single mutant males are fertile, we identified seven transcripts in *pi9^−/−^* and sixteen in *pi17^−/−^* whose steady-state abundance was significantly altered in primary spermatocytes compared to C57BL/6 controls (false discovery rate [FDR] <0.01; Figures 2A and 2B; Table S2). Among the seven dysregulated mRNAs in *pi9^−/−^* primary spermatocytes, one decreased 2.5-fold and six increased 1.5–2.4-fold (Figure 2A, at left). Five of the six derepressed mRNAs are the targets of *pi9* piRNAs (Figure 2A, at right): for each, the corresponding 5′- monophosphate-bearing 3′ cleavage product was detectable in C57BL/6 and *pi17^−/−^* and reduced ≥8-fold in *pi9^−/−^* and *pi9^−/−^pi17^−/−^* primary spermatocytes (Figure S2A).

**Figure 2.**
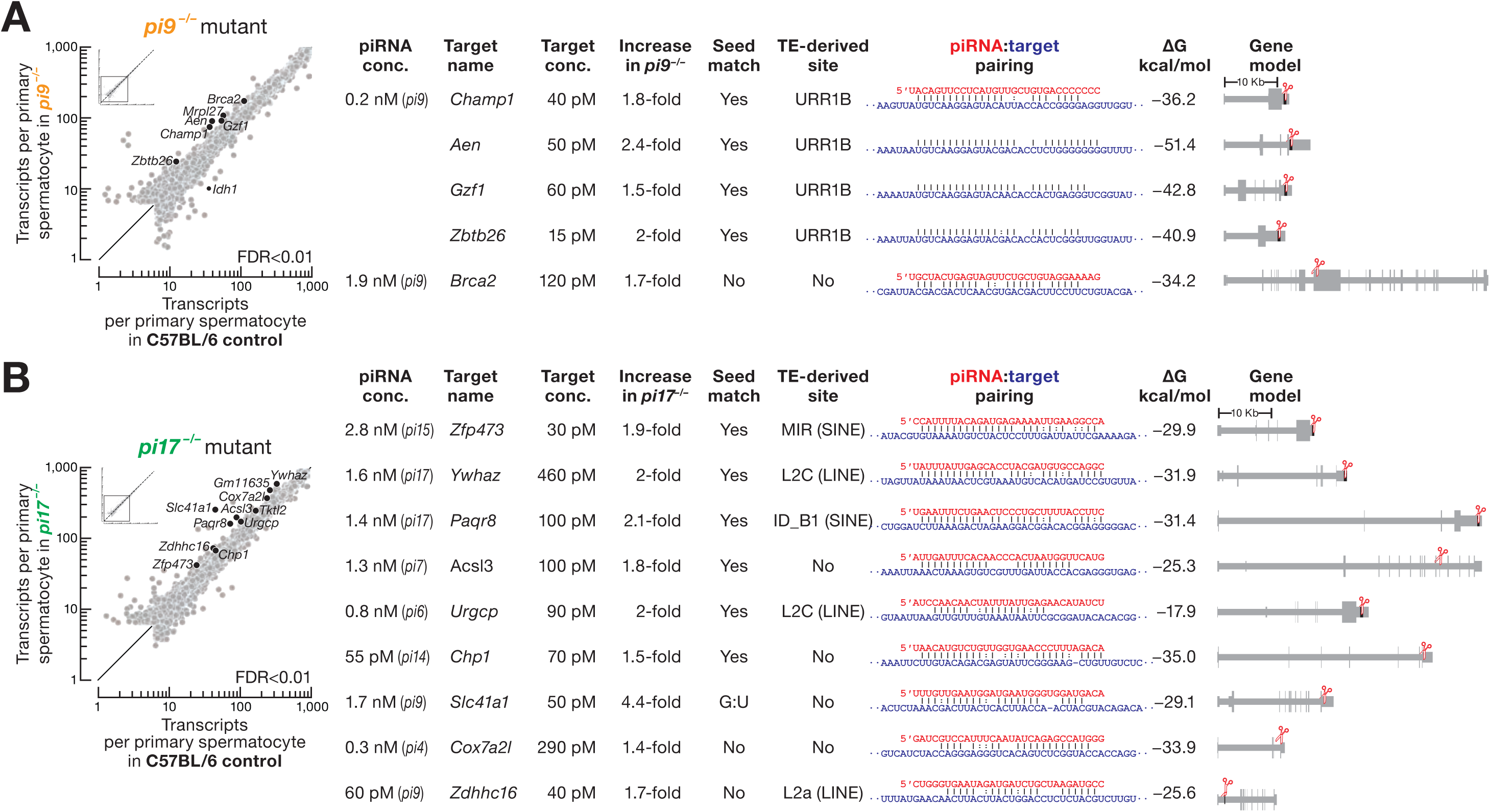
Most mRNAs Derepressed in *pi9^−/−^* and *pi17^−/−^* in Primary Spermatocytes Are Direct Targets of Pachytene piRNAs. (A) At left, scatter plot of mean steady-state transcript abundance in FACS-sorted primary spermatocytes of C57BL/6 (*n* = 7) and *pi9^−/−^*(*n* = 6) males. Differentially expressed transcripts (FDR<0.01) were identified using DESeq2 (see also STAR Methods) and are indicated. At right, direct targets of *pi9* piRNAs. piRNA and target concentrations in primary spermatocytes, extent of target increase in *pi9^−/−^* primary spermatocytes, target site location in transcript, piRNA:target pairing pattern and binding energy (computationally predicted Gibbs free energy, Δ*G*^0^) are shown. Presence of seed match (g2–g8) and whether cleavage site is found in a transposon fragment is indicated. (B) At left, scatter plot of mean steady-state transcript abundance in FACS-sorted primary spermatocytes of C57BL/6 (*n* = 7) and *pi17^−/−^*(*n* = 6) males. Differentially expressed transcripts (FDR<0.01) are shown in black. At right, direct targets of *pi17* piRNAs and piRNAs whose biosynthesis depends on *pi17*-piRNA directed cleavage. piRNA locus of origin, piRNA and target concentrations in primary spermatocytes, extent of target increase in *pi17^−/−^* primary spermatocytes, target site location in transcript, piRNA:target pairing pattern and binding energy (computationally predicted Gibbs free energy, Δ*G*^0^) are shown. Presence of seed match (g2–g8) and whether cleavage site is found in a transposon fragment is indicated. *pi4*, *4-qC5-17839*; *pi7*, *7- qD2-24830*; *pi15*, *15-qD1-17920*; *pi6*, *6-qF3-8009*; *pi14*, *14-qA3-3095*.

When *pi17* piRNAs were removed, the steady-state abundance of eleven mRNAs and five lncRNAs increased 1.4–4.4-fold; no transcripts decreased (FDR <0.01; Figure 2B, at left). Of these eleven mRNAs, our analysis suggests that four are the targets of *pi17* piRNAs and five are the targets of piRNAs whose biosynthesis is initiated by cleavage directed by a *pi17* piRNA (Figure 2B, at right). For all nine targets, we detected cleavage products in C57BL/6 and *pi9^—/—^*, and the abundance of cleavage products was reduced ≥8-fold in *pi17^−/−^*and *pi9^−/−^pi17^−/−^* primary spermatocytes (Figure S2B).

*pi9^−/−^pi17^−/−^* testes showed an elevated incidence of double-stranded DNA breaks in testicular sperm, compared to the control C57BL/6 males (40 ± 10% of tubules/section for *pi9^−/−^pi17^−/−^*vs 10 ± 9% of tubules for the control animals; two-tailed *t*-test, Welch-corrected *p*-value = 0.049: Figures 3A and 3B). Remarkably, eight of the 14 *pi9* and *pi17* target mRNAs encode proteins implicated in the DNA damage response (*Brca2*), cell proliferation (*Gzf1*, *Ywhaz*, *Acsl3*, *Zdhhc16*, *Cox7a2l, Urgcp*), or apoptosis (*Aen*; Table S3).

**Figure 3.**
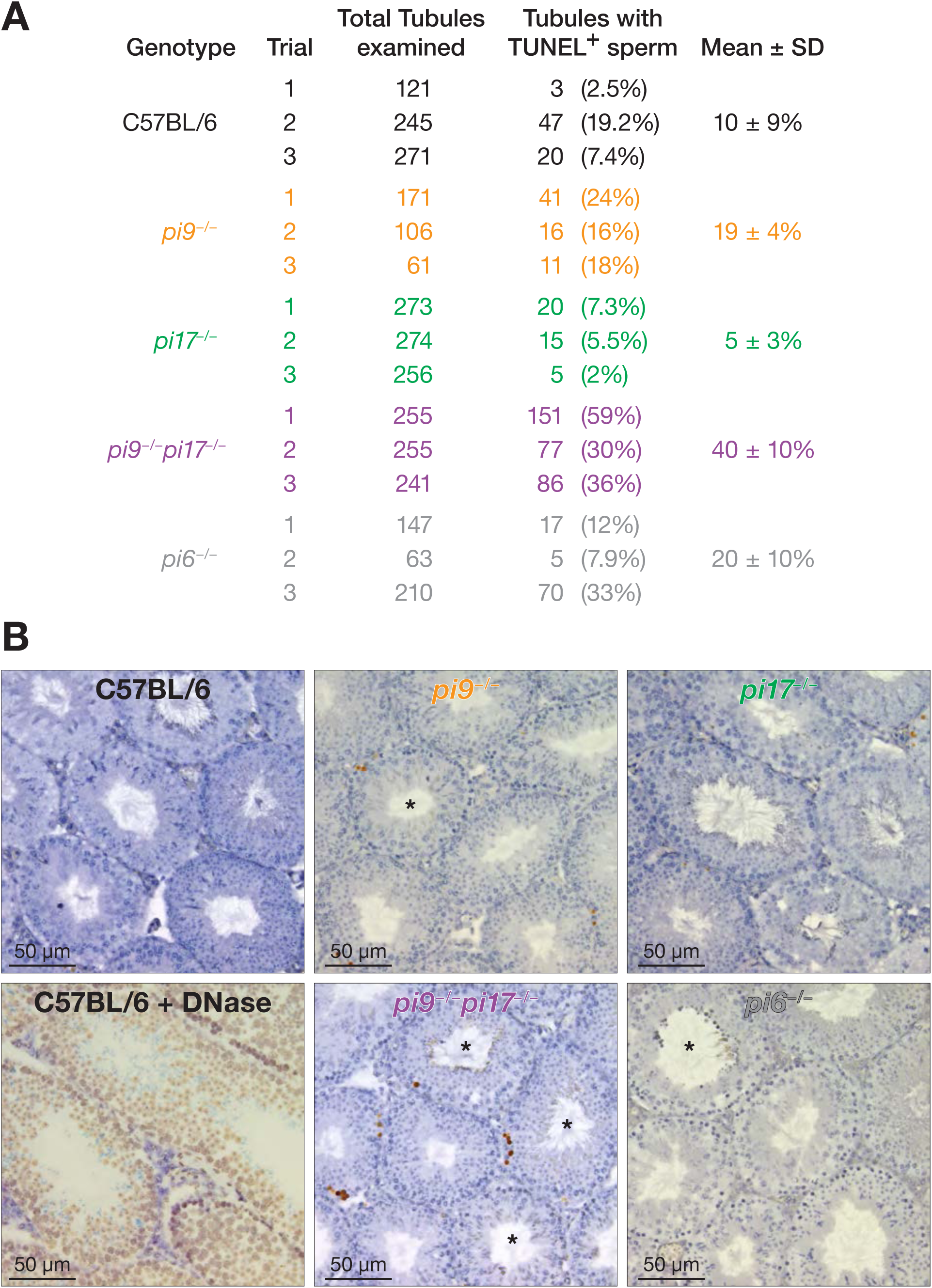
High Incidence of Double-Stranded DNA Breaks in *pi9^−/−^pi17^−/−^* Testicular Sperm. (A) Fraction of seminiferous tubules with sperm staining positive for TUNEL (terminal deoxynucleotidyl transferase dUTP nick end labeling) of C57BL/6 and pachytene piRNA mutant testes (*n* = 3 mice). (B) Representative images of seminiferous tubules of C57BL/6, C57BL/6 sections treated with DNase, and pachytene piRNA mutant testes. Brown, positive TUNEL reaction; Blue, hematoxylin.

The genomic instability observed in *pi9^−/−^pi17^−/−^*testicular sperm is consistent with the previously reported consequences of supra-physiological levels of the proteins encoded by these target mRNAs. Overexpression of a RAD51-interacting domain of BRCA2 has been shown to disrupt DNA repair by homologous recombination in human cultured cells.^31,32^ Successful meiosis requires suppression of S phase during the second round of cell division.^33^ Notably, overexpression of GZF1^34^, YWHAZ^35^, ACSL3^36^, ZDHHC16^37^, or COX7A2L^38^ stimulates mitosis in human immortalized cells and mouse cancer models. Increased abundance of URGCP has been shown stimulate the G1/S transition.^39^ Finally, an increased level of AEN (apoptosis enhancing nuclease) induces apoptosis and DNA fragmentation.^40,41^

In contrast, the incidence of dsDNA breaks observed in *pi6^−/−^*males was comparable to that in C57BL/6 controls (20 ± 10% of tubules for *p6^—/—^*vs 10 ± 9% for the controls; two-tailed *t*-test, Welch-corrected *p*-value = 0.45: Figure 2C). None of the eight mRNAs whose steady-state abundance increases in *pi6^−/−^*spermatocytes encode proteins expected to promote double-stranded DNA breaks.^20^

### Pachytene piRNAs Do Not Use a miRNA-like Mechanism to Regulate mRNA Abundance

Pachytene piRNAs were proposed to bind mRNAs via seed complementarity (piRNA nucleotides g2–g8) and recruit deadenylase CAF1 to destabilize the targets, just like miRNAs.^25^ For a single miRNA, ∼800 (median = 815, inter-quartile range [IQR] 210– 1286) high-affinity binding sites (g2–g8·t1A) are present in the UTRs of transcripts expressed in primary spermatocytes. Thus, miRNAs must be present at thousands of molecules per cell to have a detectable impact on hundreds of target mRNAs^42^: e.g., 2,500 molecules of let-7a and 23,500 molecules of miR-34c per primary spermatocyte.^19^

For *pi2*, *pi7*, *pi9*, and *pi17* pachytene piRNA loci, each gene produces between 30 and 58 piRNAs present at >1000 molecules per primary spermatocyte (Table S4). Yet loss of these piRNAs has no detectable effect on male fertility (Figure 1A), and the steady-state levels of only six mRNAs in *pi9^−/−^* and 11 mRNAs in *pi17^−/−^*change in single-mutant primary spermatocytes (Figures 2A and 2B). Conversely, loss of *pi6* piRNAs—among which there are 25 piRNAs present at >1000 molecules per cell— increases the steady-state abundance of just eight mRNAs in primary spermatocytes but causes nearly complete male sterility (Figure 1A and ref.^20^). These data are at odds with the idea that pachytene piRNAs regulate multiple targets via 7-nt seed complementarity.

Moreover, mRNAs with seed complementarity to abundant (>1000 molecules per cell) *pi9* and *pi17* piRNAs were not derepressed in *pi9^−/−^pi17^−/−^*primary spermatocytes, secondary spermatocytes, round or elongating spermatids (Figures 4A and S3A). Similarly, when we repeated these analyses for all *pi9* and *pi17* piRNAs regardless of abundance, no derepression was detected for mRNAs that paired to the piRNA seed (g2–g8) and additional ≥7 nucleotides between g9 and g30 (Figures 4A and S3A). Finally, the *Grk4* mRNA has been reported to be destabilized by a *pi17* piRNA^25^, yet its steady-state level did not change in *pi9^−/−^pi17^−/−^*males (Figure S3B). Taken together, our data argue that pachytene piRNAs do not regulate mRNA expression by a miRNA-like mechanism.

**Figure 4.**
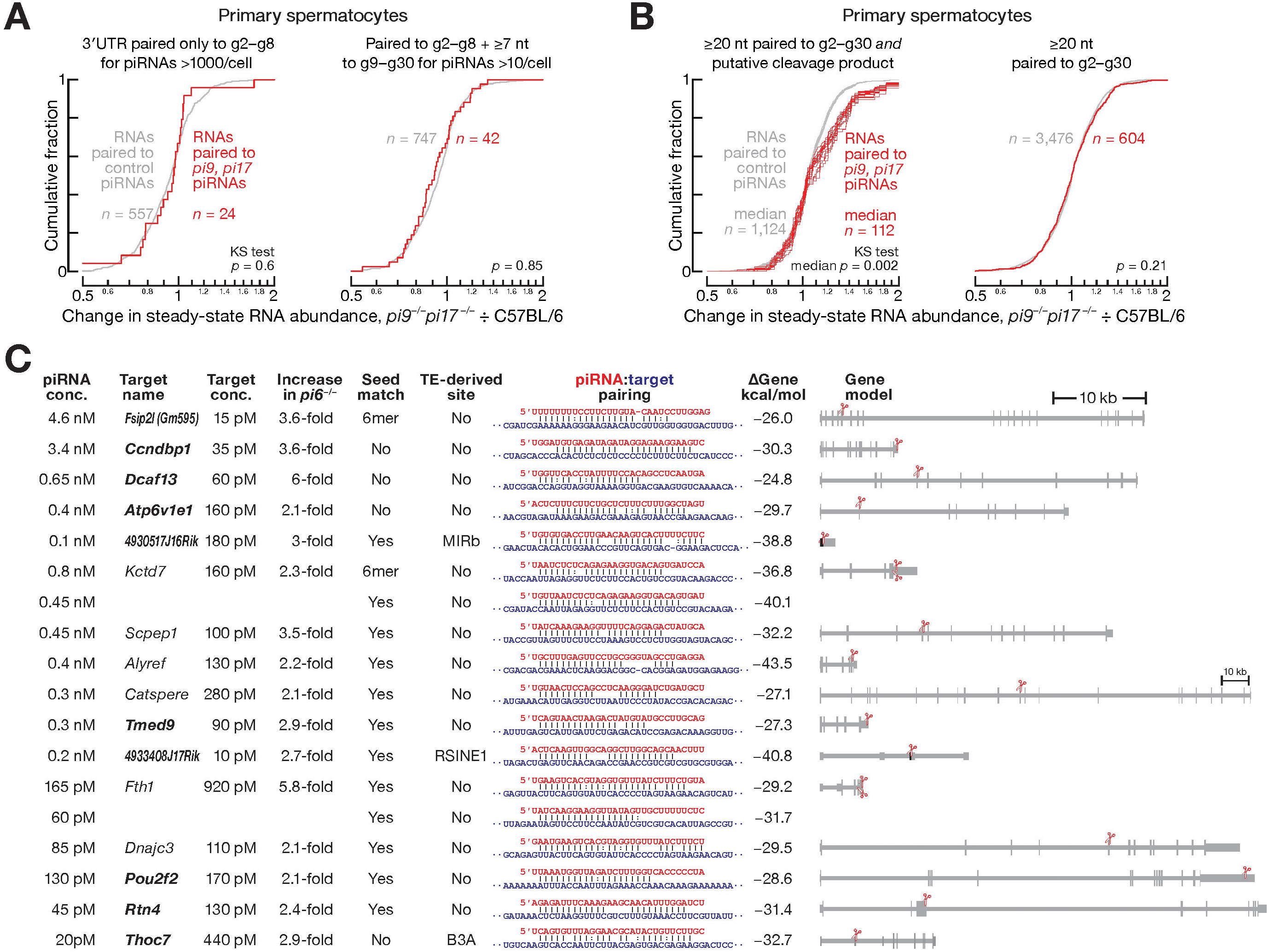
Pachytene piRNAs Regulate Targets by Endonucleolytic Cleavage. (A) Change in steady-state abundance in *pi9^−/−^pi17^−/−^*vs C57BL/6 primary spermatocytes for mRNAs whose 3′UTRs pair to *pi9* and *pi17* piRNAs or to control piRNAs (i.e. piRNAs whose abundance does not change in *pi9^−/−^pi17^−/−^* primary spermatocytes). Two-tailed Kolmogorov-Smirnov (KS) test *p*-values are shown. (B) Change in steady-state abundance in *pi9^−/−^pi17^−/−^*vs C57BL/6 primary spermatocytes for transcripts that pair with ≥20-nt to *pi9* and *pi17* piRNAs or to control piRNAs (i.e. piRNAs whose abundance does not change in *pi9^−/−^pi17^−/−^* primary spermatocytes). At left, data are only for transcripts with detectable cleavage products; *n* = 16 permutations of four control and four *pi9^−/−^pi17^−/−^*replicates of 5′- monophosphorylated RNAs. Two-tailed Kolmogorov-Smirnov (KS) test *p*-values are shown. (C) Direct targets of *pi6* piRNAs. Median target increase in *pi6^−/−^* primary spermatocytes, secondary spermatocytes, and spermatids, piRNA and target concentrations in primary spermatocytes, target site location in transcript, piRNA:target pairing pattern and binding energy (computationally predicted Gibbs free energy, Δ*G*^0^) are shown. Presence of seed match (g2–g8) and whether cleavage site is found in a transposon fragment is indicated. Cleavage of *Atp6v1e1* mRNA is guided by a *1- qC1.3-637* piRNA, whose biogenesis is initiated by *pi6* piRNA-directed cleavage.

### Pachytene piRNAs Regulate Targets by Cleavage

piRNAs direct PIWI proteins to cleave extensively complementary transposon transcripts, mRNAs, and lncRNAs in mice.^20,24,26,27,43^ In contrast to AGO-clade Argonautes, PIWI proteins readily cleave target RNAs bearing mismatches flanking the scissile bond or lacking seed complementarity.^21^ To identify pachytene piRNA-directed cleavage of mRNAs and lncRNAs, we sequenced RNAs bearing a 5′-monophosphate (i.e., a 3′ cleavage product), the hallmark of Argonaute protein-catalyzed RNA slicing. For a 5′-monophosphorylated RNA to be considered a potential target, we searched for *pi9* or *pi17* piRNAs with sufficient complementarity and intracellular abundance to have directed cleavage at that site in the full-length transcript (see STAR Methods and ref.^21^). High-confidence piRNA-target pairs were required to satisfy two additional criteria: (1) no piRNA from a locus other than *pi9* or *pi17* was predicted to have sufficient complementarity or abundance to direct cleavage of that target; and (2) the putative 3′ cleavage product was detected in the C57BL/6 controls but not in *pi9^−/−^pi17^−/−^* primary spermatocytes.

A median of 112 putative *pi9* and *pi17* targets met these requirements (IQR 86– 123 targets for 16 permutations of four control and four *pi9^−/−^pi17^−/−^* replicates of 5′- monophosphorylated RNAs; Table S5A). For comparison, we identified putative targets of piRNAs that are (1) not from *pi9* or *pi17* and (2) whose biogenesis does not depend on *pi9* or *pi17* piRNAs. For these control targets, we required (1) the non-*pi9* and non-*pi17* piRNAs to have sufficient target complementarity and abundance to be predicted to direct cleavage^21^; and (2) the putative 3′ cleavage product to be detected in both the C57BL/6 control and *pi9^−/−^pi17^−/−^* primary spermatocytes (median of 1,124 control targets; IQR 1,071–1,182; 16 permutations of 5′-monophosphorylated RNA data; Table S5B). Compared to control targets, the steady-state levels of *pi9* and *pi17* target transcripts increased significantly in *pi9^−/−^pi17^−/−^* primary spermatocytes (two-tailed Kolmogorov-Smirnov [KS] test median *p*-value *=* 0.002; IQR 0.001–0.004; Figure 4B, at left), suggesting that pachytene piRNAs regulate target levels by cleavage.

In contrast, when we selected putative targets without the requirement for a detectable 3′ cleavage product, we observed no difference in the change in steady-state levels between the control and *pi9* or *pi17* targets (two-tailed KS test *p*-value *=* 0.21; Figure 4B, at right). Collectively, these data support the idea that pachytene piRNAs regulate gene expression via cleavage of extensively complementary transcripts.

### Target Cleavage by Abundant, Partially Complementary Pachytene piRNAs Explains Most De-Repressed mRNAs in *pi6^−/−^* Mutants

The *pi6^−/−^* mutation removes >4,000 piRNAs yet increases the steady-state abundance of just 24 mRNAs in primary spermatocytes, secondary spermatocytes, and spermatids. Of these, the altered abundance of just six mRNAs was explained by cleavage directed by a *pi6* piRNA (Figure 4C; ref.^20^).

We re-examined the *pi6* data using the recently discovered piRNA targeting requirements for extent of complementarity and piRNA abundance^21^ in order to identify additional regulatory targets of *pi6* piRNAs. Because 5′-monophosphate-bearing cleavage products are short-lived in cells^44^, we sequenced 5′ monophosphorylated RNAs from control C57BL/6 and mutant *pi6^−/−^*primary spermatocytes 20-fold more deeply than previously.^20^ For *pi6* targets, the putative cleavage products were required to be detectable in C57BL/6 and reduced in abundance ≥8-fold in *pi6^−/−^* primary spermatocytes (Figure S4). These analyses identified additional ten piRNAs that explain the increased abundance of eight mRNAs and two lncRNAs in *pi6^−/−^* spermatocytes and spermatids (Figure 4C). Of the ten newly identified *pi6* piRNA-target pairs, five transcripts contained ≥1 nucleotide unpaired to the piRNA seed and three targets had mismatches at one of the nucleotides flanking the scissile phosphate (Figure 4C).

The remaining mRNAs whose derepression in *pi6^−/−^* mutants is not explained by cleavage directed by a piRNA from *pi6* had a median steady-state abundance of 10 mRNAs per primary spermatocyte (Table S2), an abundance which likely precluded detection of short-lived cleavage products derived from these transcripts. We note that in the absence of a 3ʹ cleavage product, we cannot search for piRNAs that might explain repression of these 10 mRNAs. Our data thus explain 14 of the 24 mRNAs derepressed in *pi6^−/−^* mutants by piRNA-directed cleavage. We conclude that main mechanism by which pachytene piRNAs regulate gene expression is PIWI-protein-catalyzed endonucleolytic cleavage.

### Transposon-Derived Pachytene piRNAs Are Overrepresented Among piRNAs that Cleave Targets

Pachytene piRNAs rarely derive from transposon sequences. Only 17% of piRNAs are made from repeat insertions in the precursor transcripts, half as frequently as expected from the fraction of transposons in pachytene piRNA precursors (32%; ref.^19^). However, transposon-derived piRNAs play an important role in initiating and facilitating pachytene piRNA production: repeat-derived piRNAs often cleave pachytene piRNA precursor transcripts and trigger biogenesis of more pachytene piRNAs.^19,20^

Remarkably, transposon-derived piRNAs are found 2.5-fold more frequently among those piRNAs that cleave mRNAs and lncRNAs, compared to piRNAs without an identifiable cleavage target (median = 41%, IQR 37–46% for pachytene piRNAs with cleavage targets vs 17% for piRNAs without detectable targets; one-sample, two-tailed *t*-test *p*-value = 0.00001; Figures 2A and 2B; Table S5A). We note that all repeat-derived pachytene piRNAs with cleavage targets are produced from insertions of inactive, evolutionarily old transposons (Figures 2A and 2B; Table S5A). Curiously, a single abundant *pi9* piRNA derived from the DNA transposon URR1B regulates four mRNAs (*Champ1*, *Aen*, *Gzf1*, and *Zbtb26*) via imperfect complementarity to an insertion of the same transposon family in the target 3′ UTRs (Figure 2A). Together, these results highlight the role of repeat-derived piRNAs in regulating non-transposon targets.

### Pachytene piRNAs Do Not Regulate mRNA Translational Efficiency

Mouse pachytene piRNAs, in collaboration with the protein ELAVL1, have been proposed to activate translation of partially complementary mRNAs in round spermatids.^28,29^ This model predicts that loss of translation-activating piRNAs will decrease ribosome occupancy on their mRNA targets. *pi9^−/−^pi17^−/−^* double mutant males provide an opportunity to test this prediction. Together, *pi9* and *pi17* produce ∼13.5% of all pachytene piRNAs, including both piRNAs derived from *pi9* or *pi17* transcripts as well as piRNAs from loci whose piRNA biogenesis is initiated by *pi9* or *pi17* piRNAs.^21^ Moreover, three *pi17*-derived piRNAs have been proposed to enhance the translation of three specific mRNAs.^28,29^

To examine regulation of mRNA translation during mouse spermatogenesis, we sequenced polyadenylated mRNAs and ribosome-protected mRNA footprints from FACS-purified primary spermatocytes, secondary spermatocytes, and round spermatids from C57BL/6 males. Translational efficiency—defined as ribosome occupancy normalized to mRNA abundance—increased ≥2-fold for 89 genes and decreased ≥1.5-fold for 46 genes in round spermatids compared to the preceding developmental stage of secondary spermatocytes. Consistent with the idea that translation is delayed for some mRNAs required for late spermatogenesis^45^, we found that genes whose translational efficiency increased in round spermatids were enriched for Gene Ontology classes attributed to late spermatogenesis or spermiogenesis: binding of sperm to zona pellucida, flagellated sperm motility, or spermatid development (Figure S5 and Table S6). In contrast, genes whose translational efficiency decreased in round spermatids were not enriched for spermatogenesis-specific biological processes (Figure S5 and Table S6).

To measure piRNA-dependent regulation of translational efficiency, we sequenced polyadenylated RNAs and ribosome footprints from FACS-purified primary spermatocytes, secondary spermatocytes, and round spermatids from *pi9^−/−^pi17^−/−^* males. We identified mRNAs whose 3′ UTRs contain both an ELAVL1-binding motif and a piRNA target site complementary to (1) the piRNA seed (nucleotides g2–g8) and (2) an additional ≥12 nucleotides within the piRNA region g9–g30.^28,29^ Using these data, we compared the translational efficiency for the targets of *pi9* and *pi17* piRNAs with that of the targets of piRNAs whose abundance was unchanged in *pi9^−/−^pi17^−/−^* mutants (control piRNAs). Surprisingly, the change in translational efficiency comparing *pi9^—/—^ pi17^—/—^* to C57BL/6 was indistinguishable between the control mRNAs and the *pi9* and *pi17* targets in primary spermatocytes (two-tailed KS test *p*-value *=* 0.64), secondary spermatocytes (*p =* 0.09), and round spermatids: (*p =* 0.37; Figure 5A). These results do not support the idea that pachytene piRNAs regulate translation of partially complementary mRNAs.

**Figure 5.**
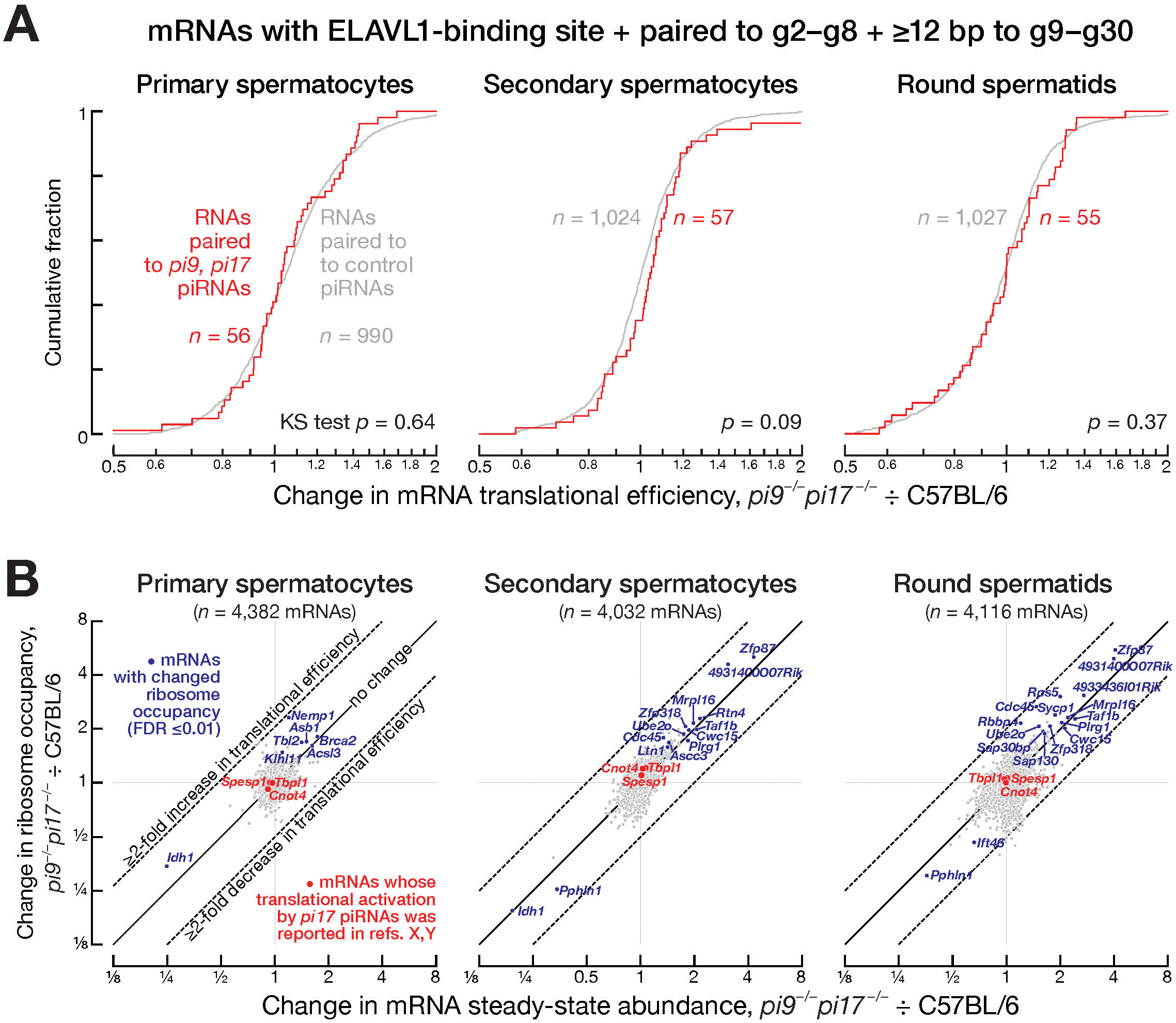
Pachytene piRNAs Do Not Regulate mRNA Translation. (A) Change in translational efficiency in FACS-purified *pi9^−/−^pi17^−/−^*vs C57BL/6 primary spermatocytes, secondary spermatocytes and round spermatids for mRNAs whose 3′UTRs pair to *pi9* and *pi17* piRNAs or to control piRNAs (i.e. piRNAs whose abundance does not change in *pi9^−/−^pi17^−/−^*germ cells). Two-tailed Kolmogorov-Smirnov (KS) test *p*-values are shown. (B) Change in steady-state abundance of mRNAs and their ribosome occupancy in FACS-purified *pi9^−/−^pi17^−/−^* vs C57BL/6 primary spermatocytes, secondary spermatocytes and round spermatids. Data are for all mRNAs with ≥10 TPM ribosome occupancy in each cell type. mRNAs with significantly changed ribosome occupancy (FDR ≤0.01) are shown in blue, mRNAs whose translational activation by *pi17* piRNAs was reported in refs.^28,29^ are shown in red.

The ribosome occupancy of only seven mRNAs in primary spermatocytes, 14 mRNAs in secondary spermatocytes, and 17 mRNAs in round spermatids changed significantly in *pi9^−/−^pi17^−/−^* mutants compared to C57BL/6 controls (FDR ≤0.01 and ≥10 TPM for ribosome occupancy in C57BL/6; Figure 5B and Table S7). We calculated the change in translational efficiency for these mRNAs in *pi9^−/−^pi17^−/−^* vs C57BL/6 by dividing the change in ribosome occupancy by the change in mRNA abundance. All changes in translational efficiency for these mRNAs were ≤2-fold (Figure 5B and Table S7). Notably, none of the 38 mRNAs with changes in ribosome occupancy were predicted to be translationally activated, because either these transcripts do not bear an ELAVL1 binding site or are not complementary to *pi9* or *pi17* piRNAs.

*Tbpl1*, *Cnot4*, and *Spesp1* have been reported to be translationally activated in round spermatids by piRNAs from *pi17*.^28,29^ We detected no change in ribosome occupancy or mRNA abundance for any of these three mRNAs in *pi9^−/−^pi17^−/−^* primary spermatocytes, secondary spermatocytes, or round spermatids (Figure 5B and Table S7). Our data do not support the proposal that pachytene piRNAs regulate mRNA translation.

### GC-2spd(ts) Cells Express Neither piRNAs Nor Most piRNA Pathway Proteins

Translational activation of *Tbpl1*, *Cnot4*, *Spesp1*, and other piRNA targets was reported to be recapitulated in GC-2spd(ts) cells, an immortalized cell line derived from mouse spermatocytes.^28,29,46,47^ Our repeated attempts to detect piRNAs in these cells were unsuccessful (Figure S6A). Nor could we detect the mRNAs encoding 15 of 20 piRNA pathway proteins in GC-2spd(ts) cells (Figure S6B), even though we could readily detect mRNAs encoding SV40 large T antigen and p53 bearing the 135 mutation, the two genes introduced in order to immortalize the original mouse pre-leptotene cells (Figure S6C and Table S8; ref.^46^). Our results corroborate an earlier report showing that GC-2spd(ts) cells do not express pre- or post-meiotic germline marker genes.^48^ We conclude that GC-2spd(ts) cells are not a useful model for pachytene piRNA biology.

### Germ Cell Granules are Unaffected by Loss of Pachytene piRNA Loci

piRNA pathway proteins localize to electron-dense structures specific to germ cells^49^, called intermitochondrial cement in primary spermatocytes and chromatoid bodies in haploid spermatids.^50–52^ In mice, intermitochondrial cement and chromatoid bodies are lost in *Miwi^−/−^* mutants: in primary spermatocytes, most mitochondria lack intermitochondrial cement, and round spermatids exhibit only diffusely staining structures presumed to be failed chromatoid bodies (Figure 6 and ref.^50^).

**Figure 6.**
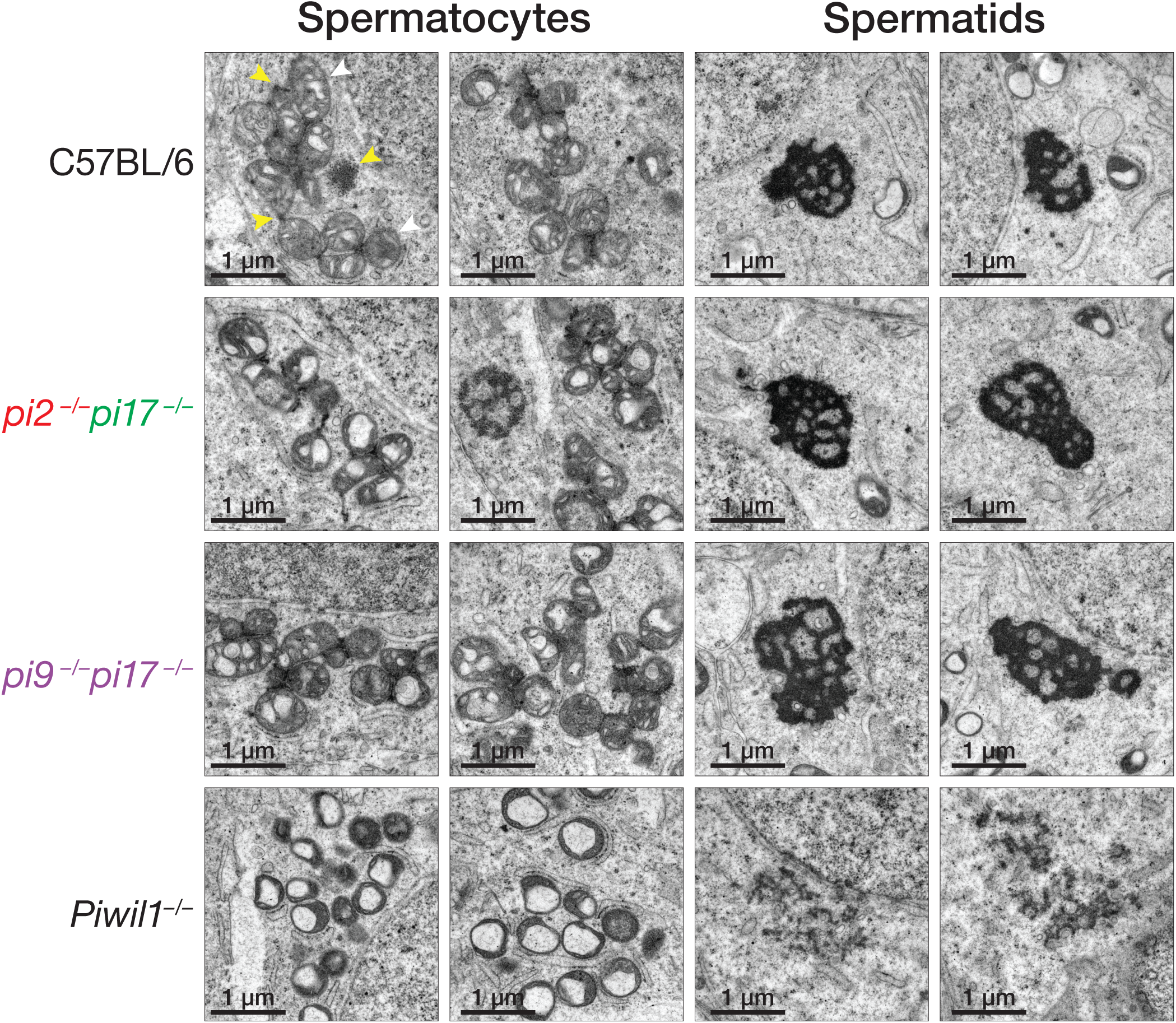
Intermitochondrial Cement and Chromatoid Body Are Intact in Pachytene piRNA Mutants. Transmission electron micrographs of spermatocytes and round spermatids from C57BL/6, *Piwil1/Miwi^−/−^*, and pachytene piRNA mutant testes.

We used transmission electron microscopy (TEM) to test whether loss of pachytene piRNA-producing loci impacts intermitochondrial cement or chromatoid bodies. The number of mitochondria and the density of intermitochondrial cement associated with them was essentially identical in primary spermatocytes of C57BL/6, *pi2^−/−^pi17^−/−^*, which are fertile but lack 12.7% of pachytene piRNAs, fertile), and *pi9^−/−^pi17^−/−^*males, which are sterile but lack 13.5% of pachytene piRNAs (Figure 6, left). Chromatoid body morphology in round spermatids from C57BL/6, *pi2^−/−^pi17^−/−^*, and *pi9^−/−^pi17^−/−^* was similarly indistinguishable (Figure 6, right). These findings demonstrate that both intermitochondrial cement and chromatoid bodies are intact in pachytene piRNA mutants, irrespective of their fertility.

### Cleavage Directed by Pachytene piRNAs Has No or Only a Modest Effect on Target Steady-State Levels

Mouse primary spermatocytes contain ∼81,600 distinct pachytene piRNA species whose concentrations range from 0.01 to 10 nM, of which *pi9* and *pi17* together produce ∼11,000 (∼13.5%; Table S4; ref.^21^). We identified a median of 112 targets cleaved by *pi9* and *pi17* piRNAs (IQR 86–123 targets for 16 permutations of four control and four *pi9^−/−^pi17^−/−^*replicates of 5′-monophosphorylated RNAs; Table S5A), suggesting that just ∼1% (i.e. 112 / 11,000) of *pi9* and *pi17* piRNAs are abundant enough and sufficiently complementary to transcripts to direct their cleavage. Consistent with high piRNA concentration being the primary determinant of efficient target cleavage^21^, the abundance of *pi9* and *pi17* piRNAs with identifiable targets (median = 123 pM; IQR 112–128 pM) was ∼3-fold higher than that of all pachytene piRNAs (38 pM; one-sample, two-tailed *t*-test *p*-value = 0.00002; Figure 7A).

**Figure 7.**
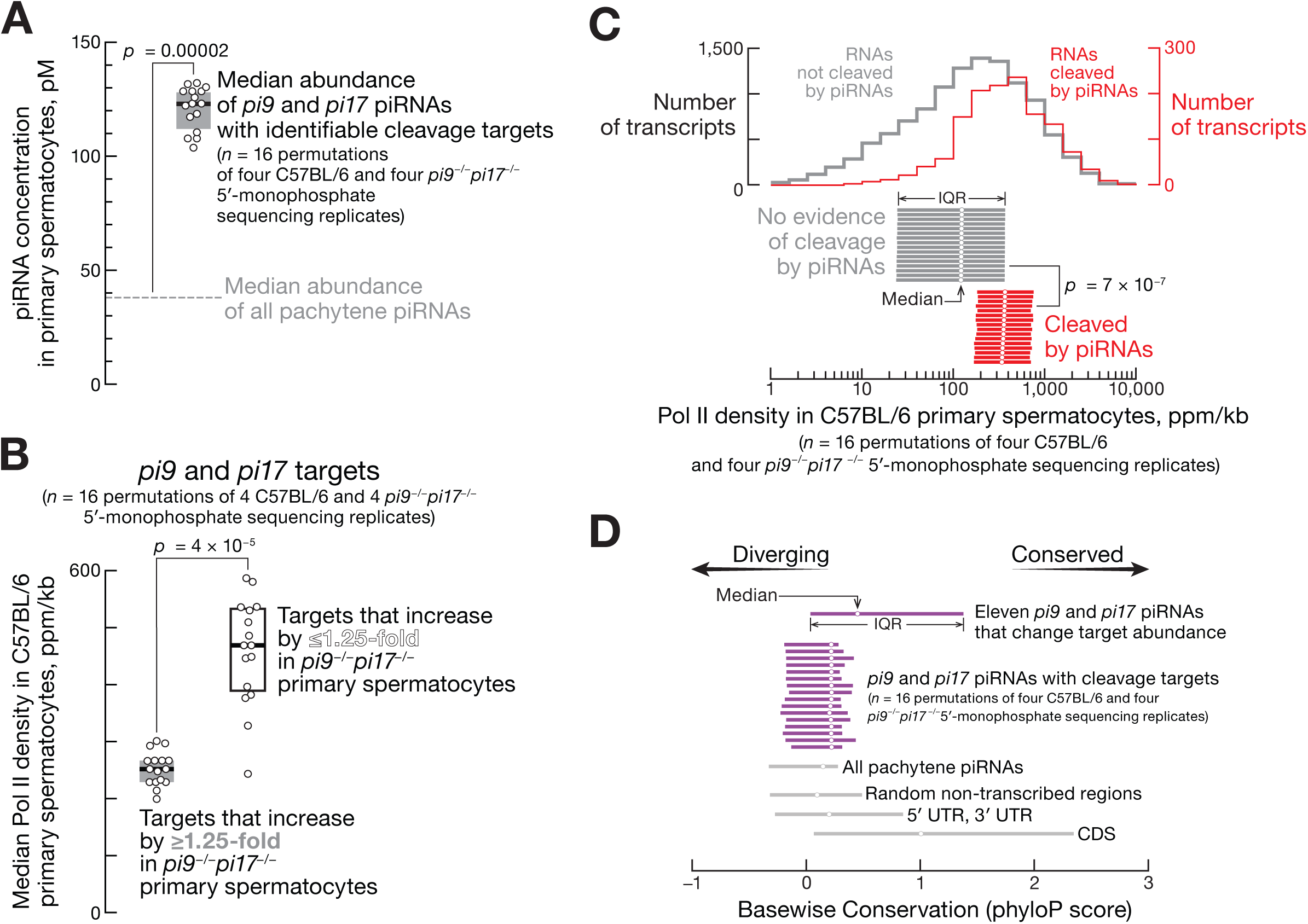
Majority of Pachytene piRNAs Target Highly Transcribed Genes. (A) Median abundance of *pi9* and *pi17* piRNAs with identifiable cleavage targets; *n* = 16 permutations of four C57BL/6 and four *pi9^−/−^pi17^−/−^* 5′-monophosphate sequencing replicates. One-sample, two-tailed *t*-test *p*-value is shown. (B) Median RNA pol II density (GRO-seq) in C57BL/6 primary spermatocytes for targets of *pi9* and *pi17* piRNAs that increase ≥1.25-fold or ≤1.25-fold in *pi9^−/−^pi17^−/−^* vs C57BL/6 primary spermatocytes; *n* = 16 permutations of four C57BL/6 and four *pi9^−/−^pi17^−/−^*5′-monophosphate sequencing replicates. Unpaired, two-tailed Mann-Whitney test *p*-value is shown. GRO-seq data are mean (*n* = 3) ppm/kb. (C) RNA pol II density in C57BL/6 primary spermatocytes for transcripts with and without detectable pachytene piRNA-directed cleavage. Top, representative distribution of RNA pol II density for one pair of C57BL/6 and *pi9^−/−^pi17^−/−^*5′- monophosphate sequencing replicates. Bottom, median and IQR for all 16 permutations of four C57BL/6 and four *pi9^−/−^pi17^−/−^* 5′-monophosphate sequencing replicates. Unpaired, two-tailed Mann-Whitney test *p*-value is shown. GRO-seq data are mean (*n* = 3) ppm/kb. (D) Base-wise conservation (PhyloP score) for genomic origins of all pachytene piRNAs, *pi9* and *pi17* piRNAs with identifiable cleavage targets (16 permutations of four C57BL/6 and four *pi9^−/−^pi17^−/−^*5′-monophosphate sequencing replicates), and eleven *pi9* and *pi17* piRNAs that change target abundance shown in Figure 2. Median and IQR for piRNA nucleotides g2–g30 are shown. For non-transcribed regions, 5′UTRs, 3′UTRs, and coding sequences, 80,000 random 29-nt long segments were sampled from each category.

Of the 112 transcripts cleaved by *pi9* and *pi17* piRNAs, a median of 71 targets (IQR 57–76) were derepressed in *pi9^−/−^pi17^−/−^* primary spermatocytes (Table S5A). For 21 (IQR 18–22) of these transcripts, the increase in steady-state level in *pi9^−/−^pi17^−/−^* primary spermatocytes met our threshold for statistical significance (FDR <0.01; Table S5A). Only eight transcripts (IQR 7–9) showed ≥1.5-fold increase in steady-state abundance in *pi9^−/−^pi17^−/−^* (Table S5A). These data suggest that most pachytene piRNA-directed cleavage events have a modest effect on target levels.

Extrapolating from the number of transcripts targeted by *pi9* and *pi17* piRNAs, we estimate that ∼830 (i.e., 112/0.135) transcripts are pachytene piRNA cleavage targets in primary spermatocytes.

### Most Pachytene piRNAs Target Highly Transcribed Genes

The combined rates of transcription and decay determine steady-state transcript abundance. Our analyses suggest that the transcription rate of most pachytene piRNA targets is higher than the rate of piRNA-directed slicing, potentially explaining why PIWI cleavage has so modest an effect on most target RNAs. To estimate the transcription rates of piRNA targets, we performed global run-on sequencing (GRO-seq) using nuclei from FACS-purified mouse primary spermatocytes. We divided the targets of *pi9* and *pi17* into two classes: those that increased ≤1.25-fold and those that increased by ≥1.25-fold in *pi9^−/−^pi17^−/−^* primary spermatocytes. Notably, median RNA pol II density for the ≤1.25-fold class was double that of the ≥1.25-fold targets (changed ≤1.25-fold: median = 470 ppm, IQR 390–530 ppm; changed ≥1.25-fold: median = 250 ppm, IQR 230–270 ppm; two-tailed, unpaired Mann-Whitney test *p*-value *=* 4 × 10^−5^; Figure 7B).

Our GRO-seq data suggest that transcripts cleaved by pachytene piRNAs generally have high transcription rates. We compared the RNA pol II density for genes with no evidence of piRNA-directed transcript slicing to that of the putative targets of all mouse pachytene piRNAs. All transcripts present in primary spermatocytes at >5 copies/cell were searched for pairing to pachytene piRNAs from the same cell type. RNAs were considered pachytene piRNA targets if (1) piRNA complementarity was predicted to be sufficient to trigger cleavage^21^ and (2) the putative cleavage product was detected in the control C67BL/6 primary spermatocytes. Transcripts bearing no or only short complementarity to pachytene piRNAs were defined as non-target controls. Remarkably, genes targeted by pachytene piRNAs exhibited ∼3-fold higher median RNA pol II signal (median = 370 ppm, IQR 330–430 ppm; two-tailed, unpaired Mann-Whitney test *p*-value *=* 7 × 10^−7^) compared to controls (median = ∼123 ppm, IQR 122– 124 ppm; Figure 7C).

Together, these data suggest that cleavage by pachytene piRNAs has a limited impact on the transcript steady-state levels due to the high transcription rate of the targets. Because pachytene piRNAs that change transcript abundance by ≥1.5-fold are rare, we hypothesize that such piRNAs are removed by purifying selection because their appearance during evolution is more frequently detrimental than advantageous for sperm fitness.

### Most *pi9* and *pi17* piRNAs That Regulate mRNAs Are Not Conserved

The promoters that initiate transcription of pachytene piRNA precursors are very similar among placental mammals.^16^ However, the primary sequences of piRNA precursor transcripts are not conserved and diverge rapidly among species.^16^

Consistent with random genetic drift in the absence of selective pressure, *pi9* and *pi17* piRNAs that cleave transcripts but do not change target steady-state levels were only marginally more conserved than piRNAs without cleavage targets (median PhyloP score for base-wise conservation for piRNAs with cleavage targets was 0.22, IQR −0.18 +0.34, vs 0.15, IQR −0.33 +0.28, for piRNAs without targets; unpaired, two-tailed Mann-Whitney test, BH-corrected *p*-value = 10^−4^;Figure 7D).

Our analyses show broad conservation of only a minority of *pi9* and *pi17* piRNAs that change mRNA steady-state levels detectably (Figure 6D). Of 14 piRNA:target pairs, the sequences of five pachytene piRNAs and their corresponding mRNA target sites are conserved among placental mammals (*Brca2*, *Slc41a1*, *Urgcp*, *Ywhaz*, *Acsl3*; Figures 2A and 2B). The other nine piRNA:target pairs are found only in the Murinae subfamily (mice and rats) of rodents. Together, these data show that even pachytene piRNAs that detectably change mRNA levels undergo rapid evolutionary turnover.

## DISCUSSION

Since the discovery of pachytene piRNAs almost two decades ago^11,12^, four models have been proposed to explain how pachytene piRNAs find and regulate their targets. Pachytene piRNAs were reported to cleave complementary transcripts, activate translation of mRNAs, destabilize mRNAs, or have no function.^20,24–30^ Our data suggest that pachytene piRNAs mainly—perhaps exclusively—regulate their targets by siRNA-like endonucleolytic cleavage.

siRNA-directed slicing by AGO proteins requires contiguous pairing from the guide 5′ seed sequence to the nucleotides past the target cleavage site.^53,54^ PIWI proteins however tolerate guide-target mismatches at any position.^21,55^ Yet even with these relaxed target finding requirements, among the tens of thousands of distinct pachytene piRNA species present in mouse primary spermatocytes at ≥10 pM, only several hundred (∼1%) are sufficiently abundant and complementary to a transcript to direct its cleavage.

Most strikingly, we find that cleavage by most pachytene piRNAs has little impact on the steady-state levels of the targets, likely due to their high transcription rates. Cleavage of the few targets whose abundance is decreased significantly by piRNAs appears to be essential for spermatogenesis. We hypothesize that pachytene piRNAs that change the steady-state levels of targets are underrepresented because such piRNAs are deleterious and removed by negative selection.

Our data thus support the idea that most pachytene piRNAs do not have a biological target^56^, because either (1) they are not complementary to any transcript or, (2) they are complementary to an RNA, but piRNA-directed cleavage does not influence the target’s steady-state abundance. These findings are consistent with the poor sequence conservation of most pachytene piRNAs.^16^ We therefore propose that the majority of pachytene piRNAs are retained in mammalian evolution because the regulation of a small number of targets by a few functional piRNAs improves spermatogenesis.

## LIMITATIONS OF THE STUDY

Detection of pachytene piRNA-guided slicing relies on sequencing of 3′ cleavage products that are 5′-monophosphorylated; because 5′-monophosphate-bearing RNAs are quickly degraded in cells, the total number of RNAs cleaved by pachytene piRNAs could be underestimated. This study does not investigate the biogenesis or function of those piRNAs (<5%) present during mouse meiosis that are not produced from the dedicated pachytene piRNA loci but are instead derived from evolutionarily young LINE1 transposons and are required for their silencing.^43^

## ACKNOWLEDGMENTS

We thank the UMass FACS Core for help sorting mouse germ cells; members of the Zamore and Gainetdinov laboratories for discussions and critical comments on the manuscript. We acknowledge the Zegar Family Foundation for their generous support and thank the NYU Center for Genomics and System Biology Genomics Core for their assistance and resources. This work was supported in part by NIGMS R35 GM136275 grant to P.D.Z., S10RR027897 to UMass Electron Microscopy Facility, and 1S10 OD028576 to UMass Flow Cytometry Core Facility. P.D.Z. is an investigator of the Howard Hughes Medical Institute.

## AUTHOR CONTRIBUTIONS

K.C., P.D.Z., and I.G. conceived the study; K.C., N.A., A.B., and I.G. performed experiments, P.D.Z. and I.G. supervised the study; K.C., P.D.Z., and I.G. wrote the manuscript.

## DECLARATION OF GENERATIVE AI AND AI-ASSISTED TECHNOLOGIES IN THE WRITING PROCESS

Generative AI and AI-assisted technologies were not used in the writing process.

## INCLUSION AND DIVERSITY

One or more of the authors of this paper self-identifies as a member of the LGBTQ+ community.

## SUPPLEMENTARY FIGURE LEGENDS

**Figure S1.**
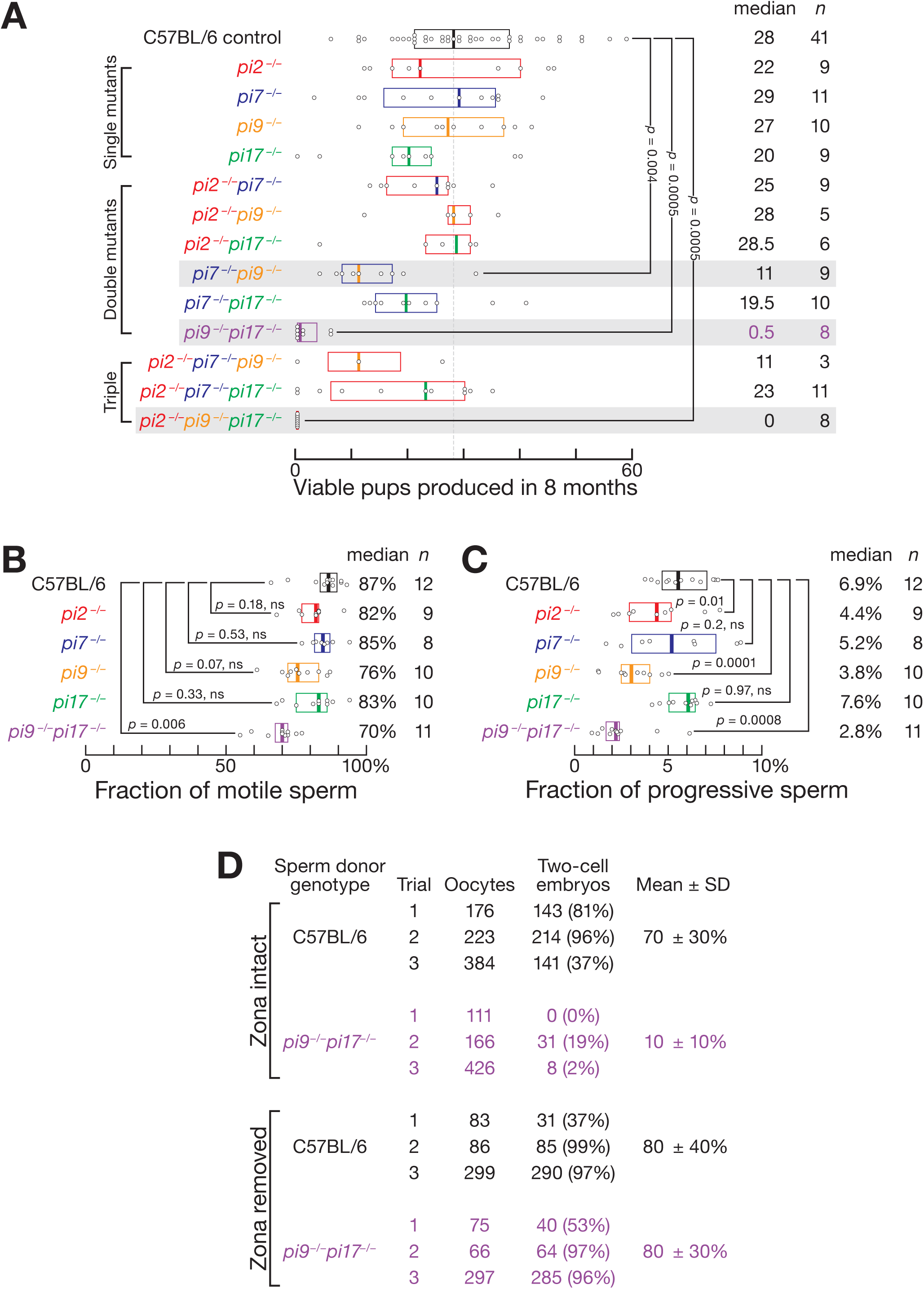
Fertility Defects Observed in Male Mice Lacking Two or Three Pachytene piRNA Loci, Related to. **Figure 1**. (A) Number of viable pups produced by the control (C57BL/6) and pachytene piRNA mutant males in successive matings with C57BL/6 females over 8 months. Median and IQR are shown. Kruskal-Wallis test (one-way ANOVA on ranks) *p*-value = 2×10^−7^. Benjamini-Hochberg corrected *p*-values for *post hoc* pairwise Mann-Whitney tests are shown. (B) Fraction of motile sperm from caudal epididymis of C57BL/6, *pi9^−/−^*, *pi17^−/−^*, and *pi9^−/−^pi17^−/−^* males determined by CASAnova. Kruskal-Wallis test (one-way ANOVA on ranks) *p*-value = 0.00036. Benjamini-Hochberg corrected *p*-values for *post hoc* pairwise Mann-Whitney tests are shown. (C) Fraction of progressive sperm from caudal epididymis of C57BL/6, *pi9^−/−^*, *pi17^−/−^*, and *pi9^−/−^pi17^−/−^* males determined by CASAnova. Kruskal-Wallis test (one-way ANOVA on ranks) *p*-value = 0.000048. Benjamini-Hochberg corrected *p*-values for *post hoc* pairwise Mann-Whitney tests are shown. (D) C57BL/6 and *pi9^−/−^pi17^−/−^* sperm function analyzed by IVF using C57BL/6 oocytes with or without zona pellucida.

**Figure S2.**
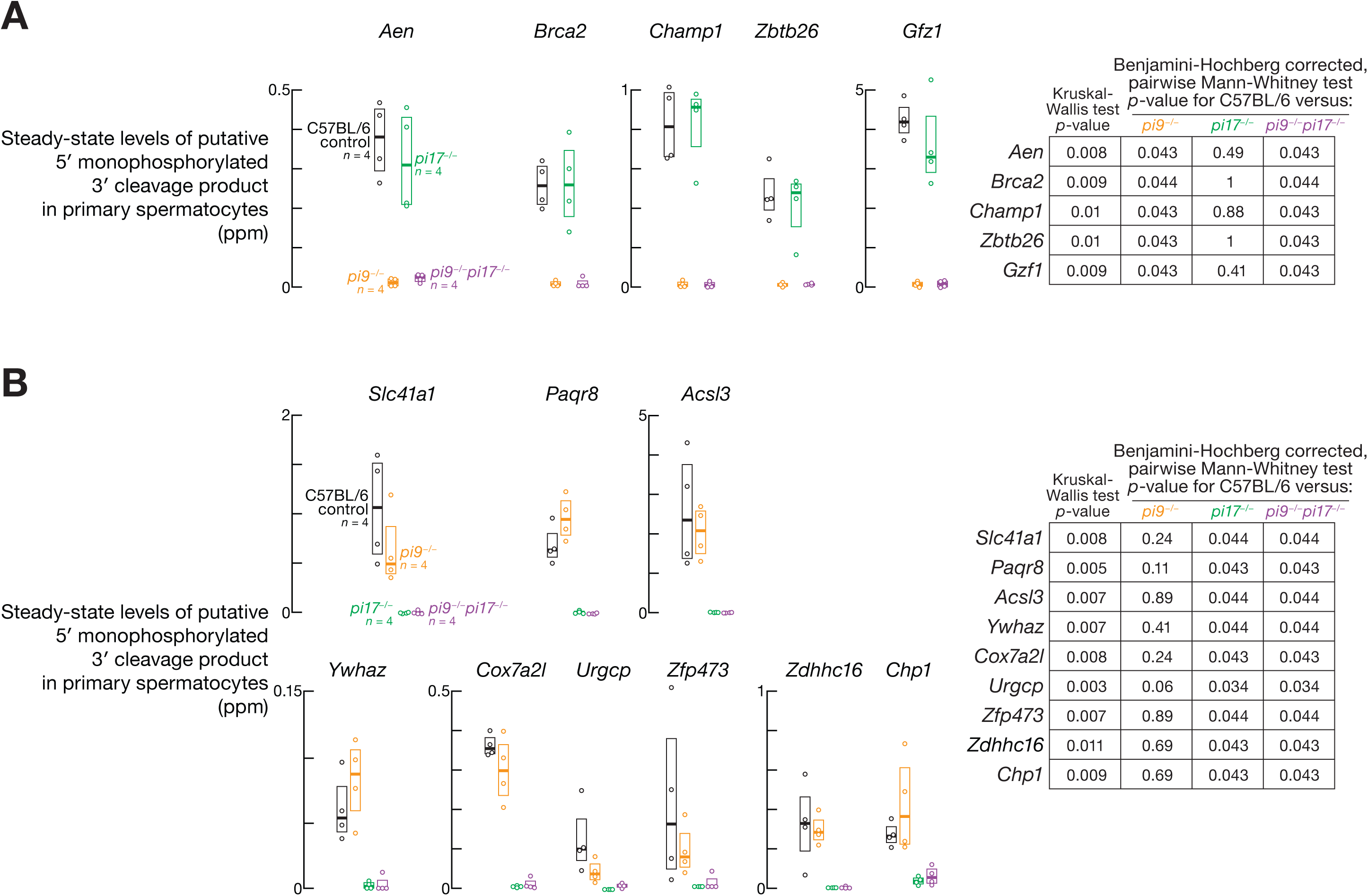
Steady-State Levels of Putative Cleavage Products Generated by *pi9* and *pi17* piRNA-Guided Slicing, Related to. **Figure 2**. (A) At left, abundance in C57BL/6, *pi9^−/−^*, *pi17^−/−^*, and *pi9^−/−^pi17^−/−^* primary spermatocytes of putative 5′-monophosphate-bearing cleavage products derived from targets of *pi9* piRNAs (shown in Figure 2A). At right, Kruskal-Wallis test *p*-values (one-way ANOVA on ranks) and Benjamini-Hochberg-corrected *p*-values for *post hoc* pairwise Mann-Whitney tests are shown for each gene. (B) At left, abundance in C57BL/6, *pi9^−/−^*, *pi17^−/−^*, and *pi9^−/−^pi17^−/−^* primary spermatocytes of putative 5′-monophosphate-bearing cleavage products derived from targets of *pi17* piRNAs (shown in Figure 2A). At right, Kruskal-Wallis test *p*-values (one-way ANOVA on ranks) and Benjamini-Hochberg-corrected *p*-values for *post hoc* pairwise Mann-Whitney tests are shown for each gene.

**Figure S3.**
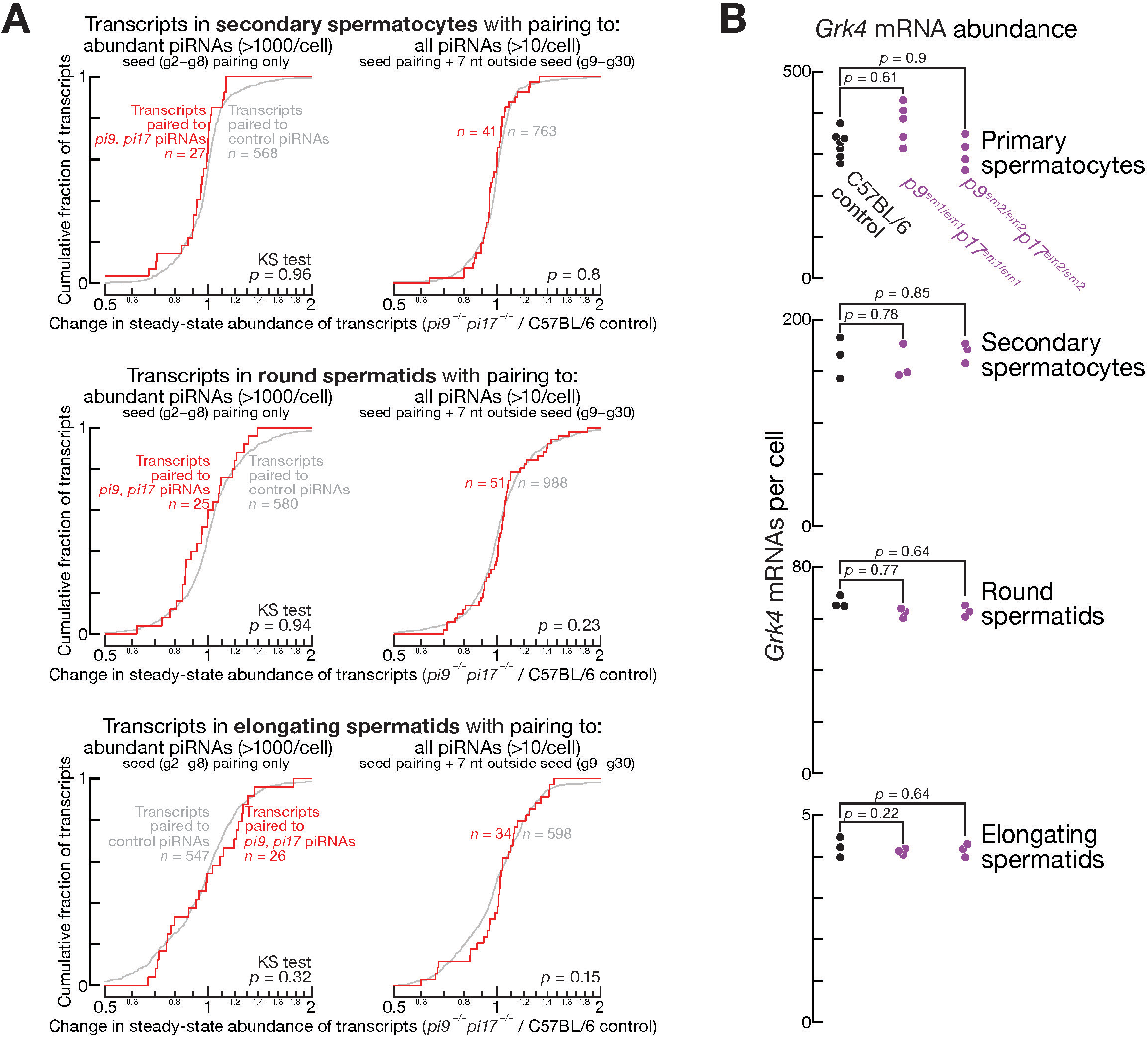
Steady-State Levels of Putative Cleavage Products Generated by *pi9* and *pi17* piRNA-Guided Slicing, Related to. **Figure 4**. (A) Change in steady-state abundance in *pi9^−/−^pi17^−/−^*vs C57BL/6 secondary spermatocytes, round spermatids, and elongating spermatids for mRNAs whose 3′UTRs pair to *pi9* and *pi17* piRNAs or to control piRNAs (i.e. piRNAs whose abundance does not change in *pi9^−/−^pi17^−/−^*primary spermatocytes). Two-tailed KS test *p*-values are shown. (B) Steady-state levels of Grk4 mRNA in C57BL/6, *pi9^em1/em1^pi17 ^em1/em1^*, and *pi9 ^em2/em2^pi17 ^em2/em2^* primary spermatocytes, secondary spermatocytes, round spermatids, and elongating spermatids. Benjamini-Hochberg-corrected *p*-values for Wald test calculated by DESeq2 are shown for each gene.

**Figure S4.**
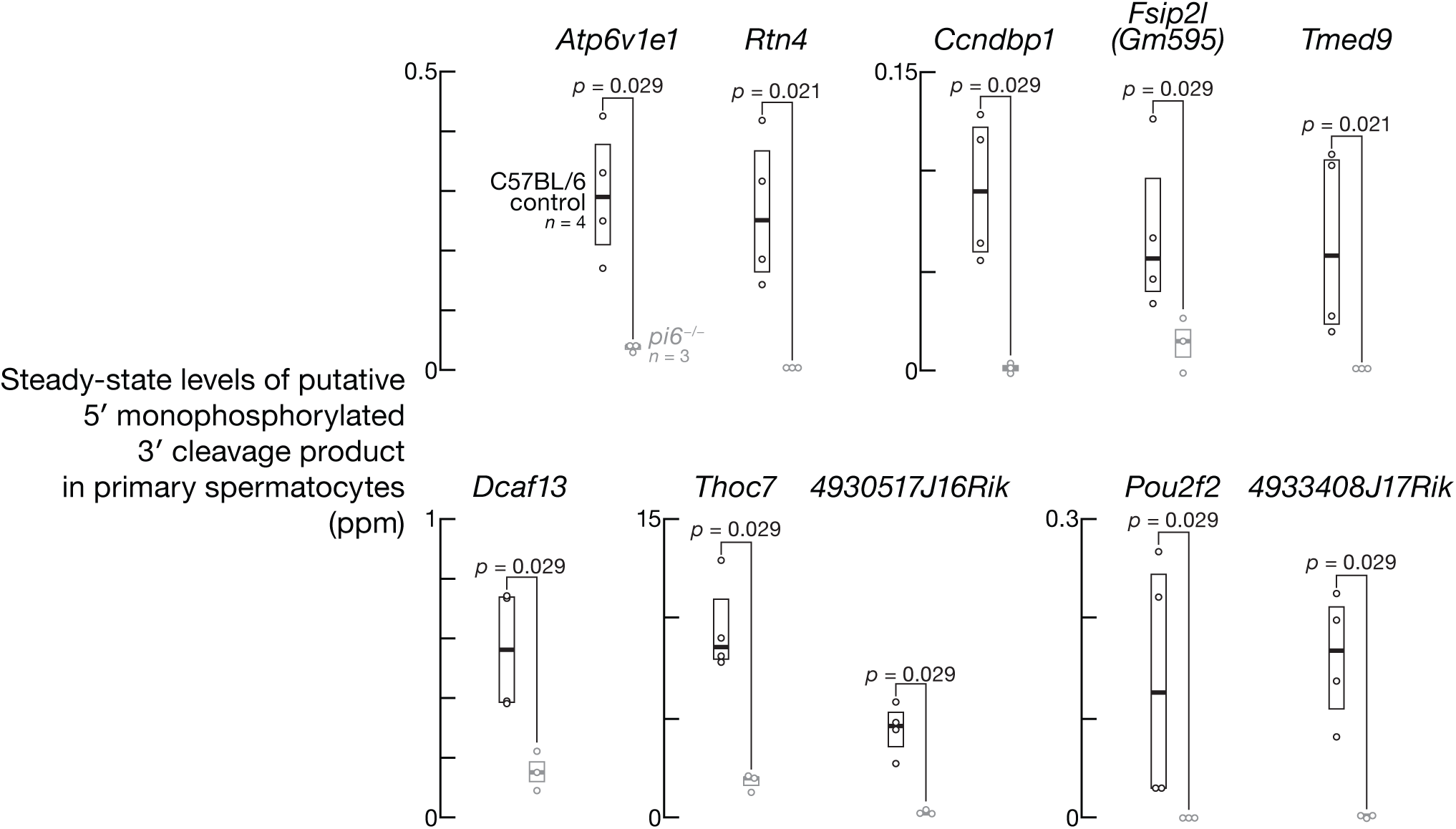
Steady-State Levels of Putative Cleavage Products Generated by *pi6* piRNA-Guided Slicing, Related to. Figure 4. Abundance in C57BL/6 and *pi6^−/−^* primary spermatocytes of putative 5′- monophosphate-bearing cleavage products derived from targets of *pi6* piRNAs (shown in Figure 4B). Unpaired, two-tailed Mann-Whitney test *p*-values are shown.

**Figure S5.**
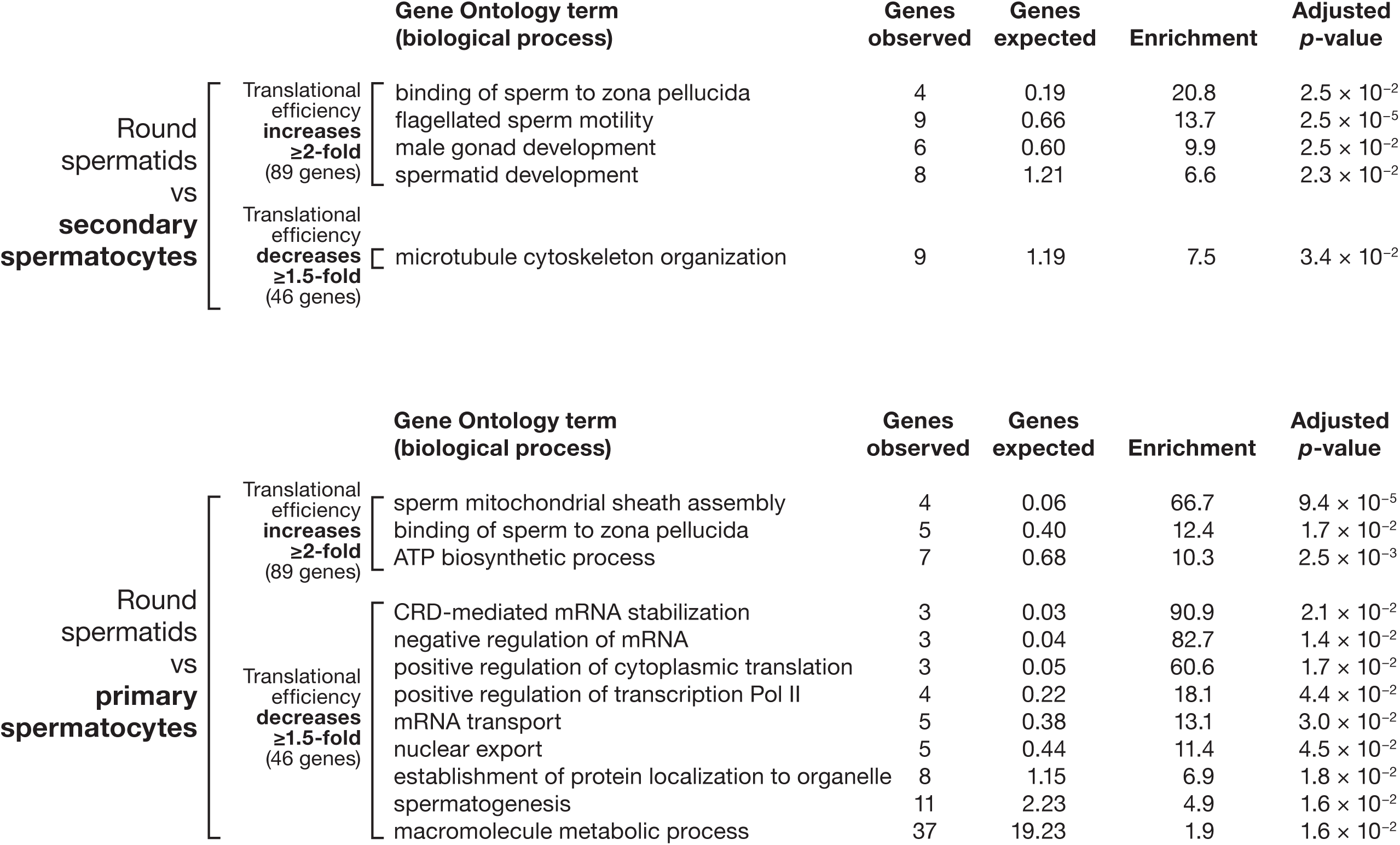
Gene Ontology Terms Enriched among mRNAs with Changed Translational Efficiency, Related to Figure 5. mRNAs whose translational efficiency increased or decreased in round spermatids compared to secondary spermatocytes (top) or primary spermatocytes (bottom) were examined for enrichment for Gene Ontology terms (biological processes) using Panther database.

**Figure S6.**
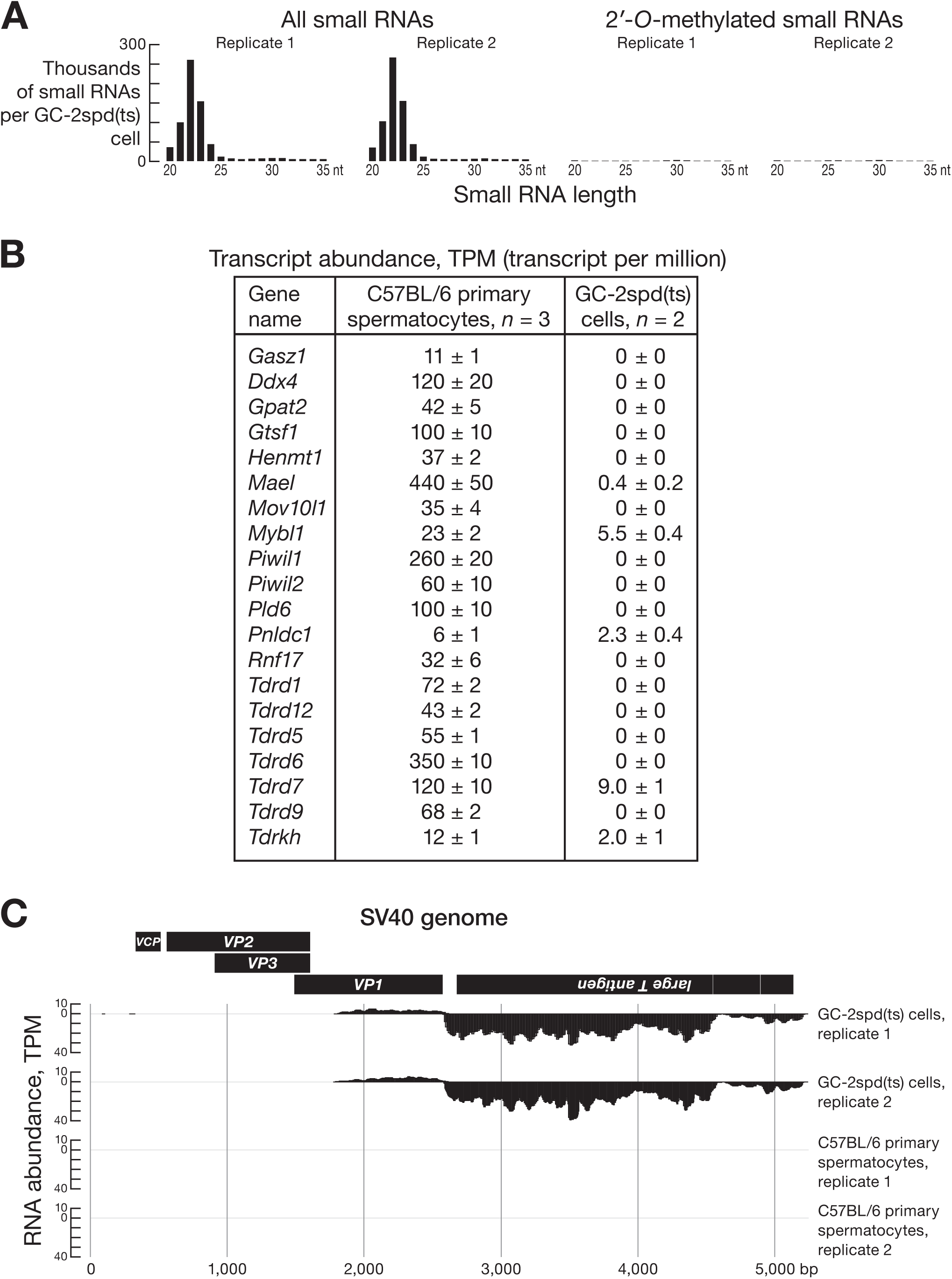
GC-2spd(ts) Cells Do Not Express piRNAs And Most piRNA Pathway Genes, Related to Figure 5. (A) Length profiles for ≥20-nt small RNAs from two biological replicates of GC-2spd(ts) cells. At left, all small RNAs were cloned and sequenced. At right, sodium periodate pre-treatment ensured cloning of only small RNAs with 2′-modified termini (e.g., 2′-*O*- methylated piRNAs). (B) Mean steady-state abundance of mRNAs encoding piRNA pathway proteins. Data are mean and SD for C57BL/6 primary spermatocytes (*n* = 3) and GC-2spd(ts) cells (*n* =2). (C) Steady-state abundance of SV40-derived transcripts in C57BL/6 primary spermatocytes (*n* = 2) and GC-2spd(ts) cells (*n* =2).

## SUPPLEMENTARY TABLES

**Table S1. Mouse strains used in this study.**

**Table S2. Change in transcript steady-state abundance in *pi9^−/−^*, *pi17^−/−^*, and *pi9^−/−^pi17^−/−^* primary spermatocytes vs C57BL/6.**

**Table S3. Known or proposed molecular function for genes whose abundance changes in *pi9^−/−^* or *pi17^−/−^* primary spermatocytes vs C57BL/6.**

**Table S4. Steady-state abundance of pachytene piRNAs in C57BL/6 primary spermatocytes.**

**Table S5. Identifiable cleavage targets of pachytene piRNAs in primary spermatocytes.**

(A) Identifiable cleavage targets of *pi9*, *pi17* piRNAs and piRNAs whose biogenesis is initiated by *pi9* or *pi17* piRNAs in primary spermatocytes.

(B) Identifiable cleavage targets of piRNAs whose abundance does not change in *pi9^−/−^pi17^−/−^* primary spermatocytes (control piRNAs).

**Table S6. Genes whose translational efficiency changes in C57BL/6 round spermatids vs secondary or primary spermatocytes.**

Data are for all RNAs with ≥10 TPM of ribosome occupancy in C57BL/6 in primary.

**Table S7. Change in steady-state abundance of poly(A)+ RNAs and ribosome footprints in *pi9^−/−^pi17^−/−^*vs C57BL/6.**

Data are for primary spermatocytes (A), primary spermatocytes (B), and round spermatids (C).

**Table S8. Sequence of *Trp53* mRNA assembled de novo using RNA-seq data from GC-2spd(ts) cells.**

## STAR METHODS

### RESOURCE AVAILABILITY

#### Lead contact

Further information and requests for resources and reagents should be directed to and will be fulfilled by the lead contacts, Phillip Zamore and Ildar Gainetdinov.

#### Materials availability

Strains generated in this study are available for non-commercial use without restriction upon request or, where indicated, from the Jackson Laboratory (https://www.jax.org/jax-mice-and-services/find-and-order-jax-mice). All other unique reagents generated in this study are available from the lead contacts.

#### Data and code availability

Sequencing data generated for this study were deposited at NCBI (https://www.ncbi.nlm.nih.gov/) and can be retrieved using accession number PRJNA1176701 (https://www.ncbi.nlm.nih.gov/bioproject/?term=PRJNA1176701). Original microscopy images have been deposited at Mendeley and are publicly available as of the date of publication. The DOI is listed in the key resources table. All other data are available in the manuscript or the supplemental information. Code used to analyze sequencing data was deposited at GitHub and can be accessed at https://github.com/ildargv/Cecchini_et_al.

### EXPERIMENTAL MODEL AND SUBJECT DETAILS

#### Mouse Strains and Mutants

Mice (wild-type C57BL/6J, IMSR # JAX:000664, RRID:IMSR_JAX:000664; *Piwil1/Miwi^−/−^*mutants, MGI: 2182488; pachytene piRNA mutants listed in Table S1) were housed and sacrificed according to the guidelines of the Institutional Animal Care and Use Committee of the University of Massachusetts Chan Medical School. Single-guide RNAs (sgRNAs; Table S1) were designed using CRISPR design tool (crispr.mit.edu/). sgRNAs were transcribed with T7 RNA polymerase and then purified by electrophoresis on 10% denaturing polyacrylamide gel. gRNA (20 ng/µl) and Cas9 mRNA (50 ng/µl, TriLink Biotechnologies, L-7206) were injected together into the pronucleus of one-cell C57BL/6 zygotes in M2 medium (Sigma, M7167). After injection, the zygotes were cultured in EmbryoMax Advanced KSOM Medium (Sigma, MR-106- D) at 37°C under 5% CO_2_ until the blastocyst stage (3.5 days), then transferred into uterus of pseudopregnant ICR females 2.5 dpc. To screen for mutant founders, genomic DNA extracted from tail tissues was analyzed by PCR using the primers listed in Table 1. All mutant strains were maintained in a C57BL/6 background; all experimental animals were the progeny of at least two backcrosses.

#### GC-2spd(ts) Cells

GC-2spd(ts) cells were purchased form ATCC (CRL-2196) and maintained in Dulbecco’s Modified Eagle’s Medium (Fisher Scientific, 11965092) containing 10% fetal bovine serum (Fisher Scientific, 10082147) at 37°C under 5% CO_2_.

## METHOD DETAILS

### Mouse fertility

Each 2–6-month-old male mouse was continuously housed with one 2–4-month-old C57BL/6 female. For male mice that did not produce pups after 3 months (∼3 cycles), the original female was replaced with a new female and the fertility test continued. To generate E8.5 or E14.5 embryos, one male mouse was housed with two C57BL/6 females. When a copulatory plug was observed, the female was housed separately until the experiment was completed.

### Epididymal sperm count

To quantify sperm abundance, cauda epididymides were collected from mice and placed in PBS. A few incisions were made in the epididymides with scissors to release the sperm, followed by incubation at 37°C and 5%CO_2_ for 20min. A 20-µl aliquot of sperm suspension was diluted in 480µl of 1% (w/v) paraformaldehyde (PFA) and sperm cells were counted using a Leica DMi8 bright-field microscope equipped with a 10×, NA 0.4 objective.

### TUNEL immunohistochemistry

Mouse testes were fixed in Bouin’s solution overnight, washed with 70% ethanol, embedded in paraffin and sectioned at 5 µm thickness. Click-iT TUNEL Colorimetric IHC Detection kit (Thermo Fisher, C10625) was used to detect DNA breaks, according to the manufacturer’s protocol. In brief, testes were fixed and embedded as described above, then were de-paraffinized in three changes of xylene for 5 min each, gradually re-hydrated in 100% (v/v), 95% (v/v), and 70% (v/v) ethanol for 5 min each, and then washed in 1× PBS for 5 min. After pre-treating the slides with 20 µg/ml Proteinase K at room temperature for 15 min, slides were washed with water twice (2 min each).

Positive control slides were treated with 1.0 U Turbo DNase (Thermo Fisher, AM2238) at room temperature for 30 min. Slides were then incubated with TdT reaction buffer containing terminal deoxynucleotidyl transferase in a humidified chamber at 37°C for 1 h. The reaction was quenched with 2x SSC for 15 min, then washed twice in PBS. Peroxidase activity was quenched in 3% (v/v) H_2_O_2_ at room temperature for 5 min. Slides were incubated with biotin azide and copper sulfate in a humidified chamber at 37°C for 30 min, then stained with peroxidase substrate at room temperature for 10 min. Nuclei were counterstained with hematoxylin I, and the slides sealed with EcoMount (Biocare Medical, EM897L). Images were captured using a Leica DMi8 bright-field microscope equipped with a 20× objective with 0.4 NA (HC PL FL L ×20/0.40 CORR PH1, Leica Microbiosystems).

### IVF and embryo transfer

IVF was performed as previously described^57^ using spermatozoa from caudal epididymis of C57BL/6 or *pi9^−/−^pi17^−/−^*mice. Spermatozoa were incubated in complete human tubal fluid media (HTF; 101.6 mM NaCl, 4.69 mM KCl, 0.37 mM KH_2_PO_4_, 0.2 mM MgSO_4_·7H_2_O, 21.4 mM Na lactate, 0.33 mM Na pyruvate, 2.78 mM glucose, 25 mM NaHCO_3_, 2.04 mM CaCl_2_·2H2O, 0.075 mg/ml penicillin-G, 0.05 mg/ml streptomycin sulfate, 0.02% (v/v) phenol red, 4 mg/ml BSA) with oocytes from B6SJLF1/J mice for 3–4 h at 37°C with constant 5% O_2_, 90% N_2_ and 5% CO_2_. Oocyte viability and the presence of pronuclei were assessed using a Nikon SMZ-2B (Nikon) dissecting microscope with a 5×, NA 0.6 objective. To observe embryo development, embryos were moved into potassium-supplemented simplex optimized media (KSOM; 95 mM NaCl, 2.5 mM KCl, 0.35 mM KH_2_PO_4_, 0.2 mM MgSO_4_·7H_2_O, 10 mM Na lactate, 0.2 mM Na pyruvate, 0.2 mM glucose, 25 mM NaHCO_3_, 1.71 mM CaCl_2_·2H_2_O, 1 mM L-glutamine, 0.01 mM EDTA, 0.075 mg/ml penicillin-G, 0.05 mg/ml streptomycin sulfate, 0.02% (v/v) phenol red, 1 mg/ml BSA; Millipore Sigma) after IVF and assessed every 24 h. To measure birth rates, two-cell embryos were transferred to Swiss Webster pseudopregnant females, and fetuses were isolated by cesarean section 18.5d after embryo transfer. For zona-free IVF, the zona pellucida of oocytes was removed with acid Tyrode’s solution as described.^58,59^

### Sperm motility

Cauda epidydimal sperm were collected from mice and placed in warm HTF media in a 37°C incubator with 5% CO_2_. A drop of sperm was removed from the suspension and pipetted into a sperm counting glass chamber, then assayed by CASA or video acquisition. CASA was conducted using an IVOS II instrument (Hamilton Thorne) with the following settings: 100 frames acquired at 60Hz; minimal contrast, 50; 4-pixel minimal cell size; minimal static contrast, 5; 0% straightness (STR) threshold; 10 μm/s VAP Cutoff; prog. min VAP, 20 μm/s; 1 0μm/s VSL Cutoff; 5-pixel cell size; cell intensity, 90; static head size, 0.30–2.69; static head intensity, 0.10–1.75; static elongation, 10–94; slow cells motile, yes; ×0.68 magnification; LED illumination intensity, 3,000; IDENT illumination intensity, 3,603; 37°C. The raw data files (that is, .dbt files for motile sperm and .dbx files for static sperm) were used for sperm motility analysis. For motile sperm, only those whose movement was captured with ≥45 consecutive frames were analyzed. For progressive/hyperactivated motility analysis, .dbt files of motile sperm were used as input for CASAnova, as previously described.^60^

### Transmission electron microscopy

Mouse caudal epididymides were dissected and immediately fixed by immersion in Karnovsky’s fixative (2% formaldehyde (v/v) and 3% glutaraldehyde (v/v) in 0.1 M sodium phosphate buffer, pH 7.4; Electron Microscopy Sciences) overnight at 4°C, and washed three times in 0.1 M phosphate buffer. Following the third wash, the tissues were postfixed in 1% osmium tetroxide (w/v; Electron Microscopy Sciences) for 1 h at room temperature, washed three more times with water for 10 min each and dehydrated using a graded series of 30%, 50%, 70%, 85%, 95%, 100% (three changes) ethanol and 100% propylene oxide (two changes) and a mixture of 50% propylene oxide (v/v) and 50% SPI-Pon 812 resin mixture (v/v; SPI Supplies). The sample was incubated in seven successive changes of SPI-Pon 812 resin over 3 d, polymerized at 68°C in flat molds and reoriented to allow cross-sectioning of spermatozoa in the lumen of epididymis. Sections measuring 70nm were cut on a Leica EM UC7 ultramicrotome (Leica Microsystems) using a diamond knife, collected on copper mesh grids and stained with 3% lead citrate (w/v) and 0.1% uranyl acetate (w/v) to increase contrast. Finally, sections were examined using a Philips CM10 transmission electron microscope (Philips Electron Optics) at 100kV. Images were recorded using the Erlangshen digital camera system (Gatan).

### FACS Isolation and Immunostaining of Mouse Germ Cells

Testes of 2–7-month-old mice were isolated, decapsulated, and incubated for 15 min at 33°C in 1× Gey′s Balanced Salt Solution (GBSS, Sigma, G9779) containing 0.4 mg/ml collagenase type 4 (Worthington, LS004188) rotating at 150 rpm. Seminiferous tubules were then washed twice with 1× GBSS and incubated for 15 min at 33°C in 1× GBSS with 0.5 mg/ml Trypsin and 1 µg/ml DNase I, rotating at 150 rpm. Next, tubules were homogenized by pipetting through a glass Pasteur pipette for 3 min at 4°C. Fetal bovine serum (FBS; 7.5% f.c., v/v) was added to inactivate trypsin, and the cell suspension was then strained through a pre-wetted 70 µm cell strainer (ThermoFisher, 22363548); cells were collected by centrifugation at 300 × *g* for 10 min. The supernatant was removed, cells resuspended in 1× GBSS containing 5% (v/v) FBS, 1 µg/ml DNase I, and 5 μg/ml Hoechst 33342 (ThermoFisher, 62249) and rotated at 150 rpm for 45 min at 33°C. Propidium iodide (0.2 μg/ml, f.c.; ThermoFisher, P3566) was added, and cells strained through a pre-wetted 40 µm cell strainer (ThermoFisher, 22363547). Spermatogonia, primary spermatocytes, secondary spermatocytes, round spermatids were purified using a FACSAria II Cell Sorter (BD Biosciences; UMass Medical School FACS Core) as described.^61,62^ Briefly, the 355-nm laser was used to excite Hoechst 33342; the 488-nm laser was used to record forward and side scatter and to excite Propidium iodide. Propidium iodide emission was detected using a 610/20 bandpass filter. Hoechst 33342 emission was recorded using 450/50 and 670/50 band pass filters. Cells were collected by centrifugation at 900 × *g* for 10 min. The supernatant was removed and cell pellets were flash frozen in liquid nitrogen and stored at −80°C.

Germ cell stages in the unsorted population and the purity of sorted fractions were assessed by immunostaining aliquots of cells. Cells were incubated for 20 min in 25 mM sucrose and then fixed on a slide with 1% (w/v) paraformaldehyde containing 0.15% (v/v) Triton X-100 for 2 h at room temperature in a humidifying chamber. Slides were washed sequentially for 10 min in: (1) PBS containing 0.4% (v/v) Photo-Flo 200 (Kodak, 1464510); (2) PBS containing 0.1% (v/v) Triton X-100; and (3) PBS containing 0.3% (w/v) BSA, 1% (v/v) donkey serum (Sigma, D9663), and 0.05% (v/v) Triton X-100. After washing, slides were incubated with primary antibodies in PBS containing 3% (w/v) BSA, 10% (v/v) donkey serum, and 0.5% (v/v) Triton X-100 overnight at room temperature in a humidified chamber. Rabbit polyclonal anti-SYCP3 (Abcam, ab15093, RRID:AB_301639, 1:1000 dilution) and mouse monoclonal anti-γH2AX (Millipore, 05- 636, RRID:AB_309864, 1:1000 dilution) were used as primary antibodies. Slides were washed again as described and then incubated with secondary donkey anti-mouse IgG (H+L) Alexa Fluor 594 (ThermoFisher, A-21203, RRID:AB_2535789, 1:2000 dilution) or donkey anti-rabbit IgG (H+L) Alexa Fluor 488 (ThermoFisher, A-21206, RRID:AB_2535792, 1:2000 dilution) for 1 h at room temperature in a humidified chamber. After incubation, slides were washed three times (10 min each) in PBS containing 0.4% (v/v) Photo-Flo 200 and once for 10 min in 0.4% (v/v) Photo-Flo 200. Finally, slides were dried and mounted in ProLong Gold Antifade Mountant with DAPI (ThermoFisher, P36931). To assess the purity of sorted fractions, 50–100 cells were staged by DNA, γH2AX, and SYCP3 staining.^61^ All samples used here met these criteria:

Spermatogonia, ∼95–100% pure with ≤ 5% pre-leptotene spermatocytes;
Primary spermatocytes, ∼10–15% leptotene/zygotene spermatocytes, ∼45–50% pachytene spermatocytes, ∼35–40% diplotene spermatocytes;
Secondary spermatocytes, ∼100%;
Round spermatids, ∼95–100%, ≤ 5% elongated spermatids.

### Small RNA-seq Library Preparation

Total RNA from sorted mouse germ cells was extracted using the mirVana miRNA isolation kit (ThermoFisher, AM1560). Small RNA libraries were constructed as described^62^ with modifications. Briefly, before library preparation, an equimolar mix of nine synthetic spike-in RNA oligonucleotides (Table S9) was added to each RNA sample to enable absolute quantification of small RNAs (Table S10). The median volume of primary spermatocytes (1,800 µm^3^) from ref.^19^ was used to calculate intracellular concentration: 1 molecule per primary spermatocyte corresponds to ∼ 1pM. To reduce ligation bias and eliminate PCR duplicates, the 3′ and 5′ adaptors both contained nine random nucleotides at their 5′ and 3′ ends, respectively (Table S9; ref.^63^) and 3′ adaptor ligation reactions contained 25% (w/v) PEG-8000 (f.c.). Briefly, 500–1,000 ng total RNA was first ligated to 25 pmol of 3′ DNA adapter (Table S9) with adenylated 5′ and dideoxycytosine-blocked 3′ ends in 30 µl of 50 mM Tris-HCl (pH 7.5), 10 mM MgCl_2_, 10 mM DTT, and 25% (w/v) PEG-8000 (NEB) with 600 U of homemade T4 Rnl2tr K227Q at 16°C overnight. After ethanol precipitation, the 50–90 nt (14–54 nt small RNA+36 nt 3′ UMI adapter) 3′ ligated product was purified from a 15% denaturing urea-polyacrylamide gel (National Diagnostics). After overnight elution in 0.4 M NaCl followed by ethanol precipitation, the 3′ ligated product was denatured in 14 µl water at 90°C for 60 s, 1 µl of 50 µM RT primer (Table S9) was added and annealed at 65°C for 5 min to suppress the formation of 5′-adapter:3′-adapter dimers during the next step. The resulting mix was then ligated to a mixed pool of equimolar amount of two 5′ RNA adapters (to increase nucleotide diversity at the 5′ end of the sequencing read, Table S9) in 20 µl of 50 mM Tris-HCl (pH 7.8), 10 mM MgCl_2_, 10 mM DTT, 1 mM ATP with 20 U of T4 RNA ligase (ThermoFisher, EL0021) at 25°C for 2 h. The ligated product was precipitated with ethanol, cDNA synthesis was performed in 20 µl at 42°C for 1 h using AMV reverse transcriptase (NEB, M0277), and 5 µl of the RT reaction was amplified in 25 µl using AccuPrime *Pfx* DNA polymerase (ThermoFisher, 12344024; 95°C for 2 min, 15 cycles of: 95°C for 15 s, 65°C for 30 s, 68°C for 15 s; primers listed in Table S9). Finally, the PCR product was purified in a 2% agarose gel. Small RNA-seq libraries samples were sequenced using a NextSeq 550 (Illumina) to obtain 79-nt, single-end reads.

### RNA-seq Library Preparation

Total RNA from sorted germ cells was extracted using the mirVana miRNA isolation kit (ThermoFisher, AM1560). RNA-seq of rRNA-depleted total RNAs was performed as described^64^ with modifications, including the addition of the ERCC spike-in mix to enable absolute quantification of RNAs and the use of unique molecular identifiers in adapters (Table S9) to eliminate PCR duplicates.^63^ Briefly, before library preparation, 1 µl of 1:100 diluted ERCC spike-in mix 1 (ThermoFisher, 4456740) was added to 1 µg total RNA. To remove rRNA, 1 µg total RNA was hybridized in 10 µl to a pool of 186 rRNA antisense oligos (0.05 µM f.c. each) in 10 mM Tris-HCl (pH 7.4), 20 mM NaCl by heating the mixture to 95°C, cooling at −0.1°C/s to 22°C, and incubating at 22°C for 5 min. RNase H (10 U; Lucigen, H39500) was added and the mixture incubated at 45°C for 30 min in 20 µl containing 50 mM Tris-HCl (pH 7.4), 100 mM NaCl, 20 mM MgCl_2_. The reaction volume was adjusted to 50 µl with 1× TURBO DNase buffer (ThermoFisher, AM2238) and then incubated with 4 U TURBO DNase (ThermoFisher, AM2238) for 20 min at 37°C. Next, RNA was purified using RNA Clean & Concentrator-5 (Zymo Research, R1016) to retain ≥ 200-nt RNAs, followed by the stranded, dUTP-based RNA-seq protocol described in ref.^64^. RNA-seq libraries were sequenced using a NextSeq 550 (Illumina) to obtain 79+79-nt, paired-end reads. The median number of all non-rRNA transcripts was ∼3,400,000 in primary spermatocytes, ∼1,700,000 in secondary spermatocytes, ∼770,000 in round spermatids and ∼50,000 in elongating spermatids. The median volume of primary spermatocytes (1,800 µm^3^) from ref.^19^ was used to calculate intracellular concentration: 1 molecule per primary spermatocyte corresponds to ∼ 1 pM.

For sequencing of polyadenylated RNAs, NEBNext Poly(A) mRNA Magnetic Isolation Module (NEB, E7490S) was used to purify poly(A)+ transcripts from 1–2 µg total RNA according to manufacturer’s instructions. Poly(A)+ RNAs were used to prepare RNA-seq libraries with NEBNext UltraExpress RNA Library Prep Kit (E3330S) except that UMI-containing adaptors (Table S9) were used. RNA-seq libraries were sequenced using an AVITI benchtop sequencer (Element Biosciences) to obtain 150+150-nt, paired-end reads.

### Sequencing of 5′ Monophosphorylated Long RNAs

Total RNA from FACS-purified primary spermatocytes was extracted using mirVana miRNA isolation kit (ThermoFisher, AM1560) and used to prepare a library of 5′ monophosphorylated long RNAs as described^19,65^ with modifications. Briefly, rRNA was depleted as described above for RNA-seq libraries. RNA was ligated to a mixed pool of equimolar amount of two 5′ RNA adapters (to increase nucleotide diversity at the 5′ end of the sequencing read, Table S9) in 20 µl of 50 mM Tris-HCl (pH 7.8), 10 mM MgCl_2_, 10 mM DTT, 1 mM ATP with 60 U of High Concentration T4 RNA ligase (NEB, M0437M) at 16°C overnight. The ligated product was isolated using RNA Clean & Concentrator-5 (Zymo Research, R1016) to retain ≥ 200-nt RNAs and reverse transcribed in 25 µl with 50 pmol RT primer (Table S9) using SuperScript III (ThermoFisher, 18080093). After purification with 50 µl Ampure XP beads (Beckman Coulter, A63880), cDNA was PCR amplified using NEBNext High-Fidelity (NEB, M0541; 98°C for 30 s; 4 cycles of: 98°C for 10 s, 59°C for 30 s, 72°C for 12 s; 6 cycles of: 98°C for 10 s, 68°C for 10 s, 72°C for 12 s; 72°C for 3 min; primers listed in Table S9). PCR products between 200–400 bp were isolated from a 1% agarose gel, purified with QIAquick Gel Extraction Kit (Qiagen, 28706), and amplified again with NEBNext High-Fidelity (NEB, M0541; 98°C for 30 s; 3 cycles of: 98°C for 10 s, 68°C for 30 s, 72°C for 14 s; 6 cycles of: 98°C for 10 s, 72°C for 14 s; 72°C for 3 min; primers listed in Table S9). The PCR product was purified from a 1% agarose gel and sequenced using a NextSeq 550 or NovaSeq (Illumina) to obtain 79+79-nt or 150+150-nt, paired-end reads.

### Sequencing of Ribosome Footprints

All steps were performed on ice, unless otherwise indicated. FACS-purified primary spermatocytes, secondary spermatocytes, or round spermatids (1–2 million cells) were lysed in 0.5 ml of 10 mM Tris-HCl (pH 7.5), 100 mM KCl, 5 mM MgCl_2_, 2 mM DTT, 1% (v/v) Triton X-100, 100 µg/ml cycloheximide (Sigma, C4859), and 1× protease inhibitor cocktail (1 mM 4-(2-Aminoethyl)benzenesulfonyl fluoride hydrochloride [Sigma; A8456], 0.3 μM Aprotinin, 40 μM betanin hydrochloride, 10 μM E-64 [Sigma; E3132], 10 μM leupeptin hemisulfate). Cell debris were removed by centrifugation at 20,000 × *g* for 10 min at 4°C. RNase I (Ambion, AM2294) was added to the supernatant (0.2 U/μl f.c.) and the sample was incubated at 25°C for 30 min and then moved to a polycarbonate ultracentrifuge tube (Beckman Coulter, 362305). A 3 ml sucrose cushion (10 mM Tris-HCl pH 7.5, 100 mM KCl, 5 mM MgCl_2_, 2 mM DTT, 100 µg/ml cycloheximide [Sigma, C4859], 20 U/ml SUPERaseIn RNase Inhibitor (Fisher Scientific, AM2694) in 1 M sucrose) was placed under the sample using a 21G needle (BD, 305167) on a 5 ml syringe (Fisher Scientific, 14955458). Ribosomes were precipitated by centrifugation at ∼400,000 × *g* for 90 min at 4°C (100,000 rpm in TLA-110 rotor in Optima MAX-XP Benchtop Ultracentrifuge). RNA was extracted from the ribosome pellet using mirVana miRNA isolation kit (ThermoFisher, AM1560). After ethanol precipitation, the 27–33-nt ribosome footprints were purified from a 15% denaturing urea-polyacrylamide gel (National Diagnostics). After overnight elution in 0.4 M NaCl followed by ethanol precipitation, the 3′ ends of ribosome footprints we dephosphorylated at 37°C for 4 h in 50 μl of 100 mM MES-NaOH (pH 5.5), 300 mM NaCl, 10 mM MgCl_2_, 1 U/μl SUPERaseIn RNase Inhibitor (Fisher Scientific, AM2694), 15 mM 2-mercaptoethanol, 0.8 U/μl T4 PNK (NEB, M0201). After ethanol precipitation, ribosome footprints were ligated to 25 pmol of 3′ DNA adapter for small RNA sequencing (Table S9) with adenylated 5′ and dideoxycytosine-blocked 3′ ends in 30 µl of 50 mM Tris-HCl (pH 7.5), 10 mM MgCl_2_, 10 mM DTT, and 25% (w/v) PEG-8000 (NEB) with 600 U of homemade T4 Rnl2tr K227Q at 16°C overnight. After ethanol precipitation, the 5′ ends of 63–69 nt 3′ ligated product (27–33-nt footprints plus 36-nt 3′ UMI adapter) were phosphorylated in 20 µl of 70 mM Tris-HCl (pH 7.6), 10 mM MgCl_2_, 5 mM DTT, 1 mM ATP with 20 U of T4 PNK (NEB, M0201). Following an ethanol precipitation, RNAs were denatured in 14 µl water at 90°C for 60 s, 1 µl of 50 µM RT primer (Table S9) was added and annealed at 65°C for 5 min to suppress the formation of 5′-adapter:3′- adapter dimers during the next step. The resulting mix was then ligated to a mixed pool of equimolar amount of two 5′ small RNA-seq adapters (to increase nucleotide diversity at the 5′ end of the sequencing read, Table S9) in 20 µl of 50 mM Tris-HCl (pH 7.8), 10 mM MgCl_2_, 10 mM DTT, 1 mM ATP with 20 U of T4 RNA ligase (ThermoFisher, EL0021) at 25°C for 2 h. The ligated product was precipitated with ethanol, cDNA synthesis was performed in 20 µl at 42°C for 1 h using AMV reverse transcriptase (NEB, M0277), and 5 µl of the RT reaction was amplified in 25 µl using AccuPrime *Pfx* DNA polymerase (ThermoFisher, 12344024; 95°C for 2 min, 16 cycles of: 95°C for 15 s, 65°C for 30 s, 68°C for 15 s; primers listed in Table S9). Finally, the PCR product was purified in a 2% agarose gel. Ribosome footprint libraries were sequenced using a NextSeq 550 (Illumina) to obtain 79-nt, single-end reads.

### GRO-seq

All steps were performed on ice, unless otherwise indicated. FACS-purified primary spermatocytes (1–3 million cells) were collected by centrifugation at 400 × *g* for 10 min at 4°C. Supernatant was removed and cells were carefully resuspended by pipetting in 1 ml swelling buffer (10 mM Tris-HCl (pH 7.5), 3 mM CaCl_2_, 2 mM MgCl_2_). Additional 9 ml of swelling buffer was added, cells were mixed by swirling and incubated on ice for 5 min. After collecting swollen cells by centrifugation at 400 × *g* for 10 min at 4°C, supernatant was removed and cells were resuspended in 500 μl of Lysis buffer: 10 mM Tris-HCl (pH 7.5), 3 mM CaCl_2_, 2 mM MgCl_2_, 10% glycerol, 0.04 U/µl RNasin PLUS (Promega, N2615), and 1× protease inhibitor cocktail (1 mM 4-(2- Aminoethyl)benzenesulfonyl fluoride hydrochloride [Sigma; A8456], 0.3 μM Aprotinin, 40 μM betanin hydrochloride, 10 μM E-64 [Sigma; E3132], 10 μM leupeptin hemisulfate). While carefully swirling the tube, 500 µl of Lysis buffer containing 1% Igepal CA-630 was added by drop and cells were lysed for 5 min on ice. Additional 9 ml of Lysis buffer containing 0.5% Igepal CA-630 was added, lysate was mixed by swirling, nuclei were collected by centrifugation at 600 × *g* for 5 min at 4°C, supernatant was removed, and nuclei were resuspended by pipetting in 1 ml of Lysis buffer containing 0.5% Igepal CA-630. Additional 9 ml of Lysis buffer containing 0.5% Igepal CA-630 was added, nuclei were mixed by swirling and collected by centrifugation at 600 × *g* for 5 min at 4°C, supernatant was removed, and nuclei were resuspended in 1 ml of Freezing buffer (50 mM Tris-HCl pH 8.0, 5 mM MgCl_2_, 5 mM EDTA, 40% glycerol, 0.4 U/µl RNasin PLUS (Promega, N2615), 1× protease inhibitor cocktail). Nuclei were collected by centrifugation at 900 × *g* for 5 min at 4°C, supernatant was removed, and nuclei were resuspended in 100 μl of Freezing buffer, flash frozen in liquid nitrogen, and stored at −80°C.

For nuclear run-on reaction, 100 μl of frozen nuclei was thawed on ice for 5 min and then mixed with 100 μl of 10 mM Tris-HCl pH 8.0, 300 mM KCl, 5 mM MgCl_2_, 10 mM DTT, 0.5 mM of each ATP, GTP, CTP, and BrdUTP (Sigma, B7166), 1% N-Lauroylsarcosine (Sigma L7414), 1 U/µl RNasin PLUS (Promega, N2615), and 1× protease inhibitor cocktail. Reaction was mixed with P200 tip with its end cut off and incubated at 30°C for 30 min, then 24 µl of 10× TURBO DNase buffer and 10 µl of TURBO DNase (2 U/µl, Fisher Scientific, AM2238) were added, reaction was incubated at 37°C for 20 min and RNA was extracted with Trizol, resuspended in 30 µl water and stored at −80°C.

To capture BrdU-labelled nascent transcripts, 30 µl of the sample from previous step was incubated at 65°C for 5 min, chilled on ice, and mixed with 270 µl of IP buffer: 50 mM Tris-HCl pH 8.0, 150 mM NaCl, 1 mM DTT, 1 mM EDTA, 0.05% Tween-20, 1 U/µl RNasin PLUS (Promega, N2615), and 1× protease inhibitor cocktail. Anti-BrdU mouse biotin-conjugated antibody (1 µg, 5 µl of 0.2 µg/µl of Clone PRB-1, MilliporeSigma, MAB3262BMI) was added to the RNAs in IP buffer and RNAs were incubated at 4°C for 1 h with rotation. In a separate tube, 50 µl of Dynabeads MyOne Streptavidin T1 (Fisher Scientific, 6560) were washed at room temperature for 5 min in 1 ml of IP buffer, and beads were then blocked at room temperature for 1 h with rotation in 300 µl of IP buffer containing 0.1% polyvinylpyrrolidone and 1 mg/ml Ultrapure BSA (Fisher Scientific, AM2618). After blocking, supernatant was removed, and beads were resuspended in solution containing RNAs and antibody from previous step. Biotin-conjugated antibody was allowed to bind streptavidin beads at 4°C for 30 min with rotation. Beads were then washed five times with IP buffer at 4°C for 5 min with rotation, and RNAs were extracted with Trizol. GRO-seq libraries were constructed using the method described for rRNA-depleted RNA-seq libraries and sequenced with a NextSeq 550 (Illumina) to obtain 79+79-nt, paired-end reads.

## QUANTIFICATION AND STATISTICAL ANALYSIS

### Analysis of Small RNA Sequencing Data

The 3′ adapter (5′-TGGAATTCTCGGGTGCCAAGG-3′) was removed with fastx toolkit (v0.0.14), PCR duplicates were eliminated as described^63^, and rRNA matching reads were removed with bowtie (parameter -v 1; v1.0.0; ref.^66^) against Mus musculus set in SILVA rRNA database^67^. Deduplicated and filtered data were analyzed with Tailor^68^ to account for non-templated tailing of small RNAs. Sequences of synthetic spike-in oligonucleotides (Table S9) were identified allowing no mismatches with bowtie (parameter -v 0; v1.0.0; ref.^66^), and the absolute abundance of small RNAs calculated (Table S10). Because piRNA 3′ trimming by PNLDC1 results in piRNA 3′ end heterogeneity, sequencing reads were next grouped by their 5′, 25-nt prefix. For further analyses, we kept only prefix groups that two three criteria. First, the prefix group total abundance was ≥ 1 ppm (i.e., ≥ 10 piRNAs/mouse primary spermatocyte), ensuring that, assuming a Poisson or a Negative Binomial distribution for piRNA concentration in different cells, ≥99.99% of primary spermatocytes contained at least one molecule of the piRNA 25-nt prefix. Second, the prefix group total abundance was ≥1 ppm in all 12 replicates of control C57BL/6 samples (Table S10). piRNAs were considered undetectable in *pi6*^−/−^, *pi9*^−/−^, *pi17*^−/−^ or *pi9*^−/−^*pi17*^−/−^ primary spermatocytes if their mean abundance in mutants was ≤ 0.1 ppm.

### Analysis of RNA-seq Data

RNA-seq analysis was performed using piPipes for genomic alignment^69^. Briefly, before starting piPipes, sequences were reformatted to extract unique molecular identifiers^63^. The reformatted reads were then aligned to rRNA using bowtie2 (v2.2.0; ref.^70^). Unaligned reads were mapped to mouse genome mm10 using STAR (v2.3.1; ref.^71^), and PCR duplicates removed.^63^ Transcript abundance was calculated using StringTie (v1.3.4; ref.^72^). Differential expression analysis was performed using DESeq2 (v1.18.1; ref.^73^). Only significant (FDR<0.01) changes in gene expression observed in both alleles for each pachytene piRNA mutant were considered (Table S2). De novo assembly of *Trp53* mRNA sequence was performed with Trinity (v2.14.0; ref.^74^).

### Analysis of 5′ Monophosphorylated Long RNA Sequencing Data

Sequencing data for 5′ monophosphorylated long RNAs was aligned to the mouse genome with piPipes^69^. Briefly, before starting piPipes, the degenerate portion of the 5′ adapter sequences were removed the (nucleotides 1–15 of read1). Because each library was sequenced at least twice to increase the sequencing depth, to harmonize the length of paired-end reads from different runs, sequences were trimmed to 64-nt (read1) + 79-nt (read2) paired reads. The trimmed reads were then aligned to rRNA using bowtie2 (v2.2.0; ref.^70^). Unaligned reads were mapped to mouse genome mm10 using STAR (v2.3.1; ref.^71^), alignments with soft clipping of ends were removed with SAMtools (v1.0.0; ref.^75^), and reads with the same 5′ end were merged to represent a single 5′ monophosphorylated RNA species. For further analyses, only unambiguously mapping 5′ monophosphorylated RNA species were used. For 5′ monophosphorylated RNAs mapped in annotated transcripts, the nucleotide sequence of the corresponding transcript was used to find piRNAs potentially explaining the cleavage, and we used the genomic sequence for 5′ monophosphorylated RNAs mapped outside any annotated transcript.

Search for putative cleavage targets of *pi9* and *pi17* piRNAs in *pi9^−/−^pi17^−/−^*primary spermatocytes (Figure 4B and Table S5) was performed with a threshold of ≥0.1 ppm for 5′ monophosphorylated putative cleavage products and the following piRNA-target pairing patterns were considered:

- for piRNAs at ≥1 ppm (10 molecules per primary spermatocyte), ≥20 nt paired between g2–g25;
- for piRNAs at ≥5 ppm (50 molecules per primary spermatocyte), contiguous pairing between g3–g15;
- for piRNAs at ≥10 ppm (100 molecules per primary spermatocyte), contiguous pairing between g3–g16;
- for piRNAs at ≥50 ppm (500 molecules per primary spermatocyte), contiguous pairing between g4–g17.

### Analysis of Ribosome Footprint Sequencing Data

The 3′ adapter (5′-TGGAATTCTCGGGTGCCAAGG-3′) was removed with fastx toolkit (v0.0.14), PCR duplicates were eliminated as described^63^, and rRNA matching reads were removed with bowtie (parameter -v 1; v1.0.0; ref.^66^) against Mus musculus set in SILVA rRNA database^67^. Unaligned reads were mapped to mouse genome mm10 using STAR (v2.3.1; ref.^71^). Ribosome occupancy was calculated using StringTie (v1.3.4; ref.^72^). Differential expression analysis was performed using DESeq2 (v1.18.1; ref.^73^). Only significant (FDR<0.01) changes in gene expression observed in both alleles for each pachytene piRNA mutant were considered (Table S2). Identification of mRNAs with ELAVL1-binding motif was as previously described in ref.^28^.

### Analysis of GRO-seq Data

GRO-seq analysis was performed using piPipes for genomic alignment^69^. Briefly, before starting piPipes, sequences were reformatted to extract unique molecular identifiers^63^. The reformatted reads were then aligned to rRNA using bowtie2 (v2.2.0; ref.^70^). Unaligned reads were mapped to mouse genome mm10 using STAR (v2.3.1; ref.^71^), and PCR duplicates removed^63^. RNA pol II density was calculated using BEDTools genomecov (v2.3.4; ref.^76^) as read coverage normalized by sequencing depth and gene length (parts per million per kb; ref.^72^). To minimize any contribution from paused RNA pol II, the first 500 bp of genes were excluded from analyses.

### Statistical Tests

*P* values were calculated using statistical tests described in figure legends.

## KEY RESOURCES TABLE

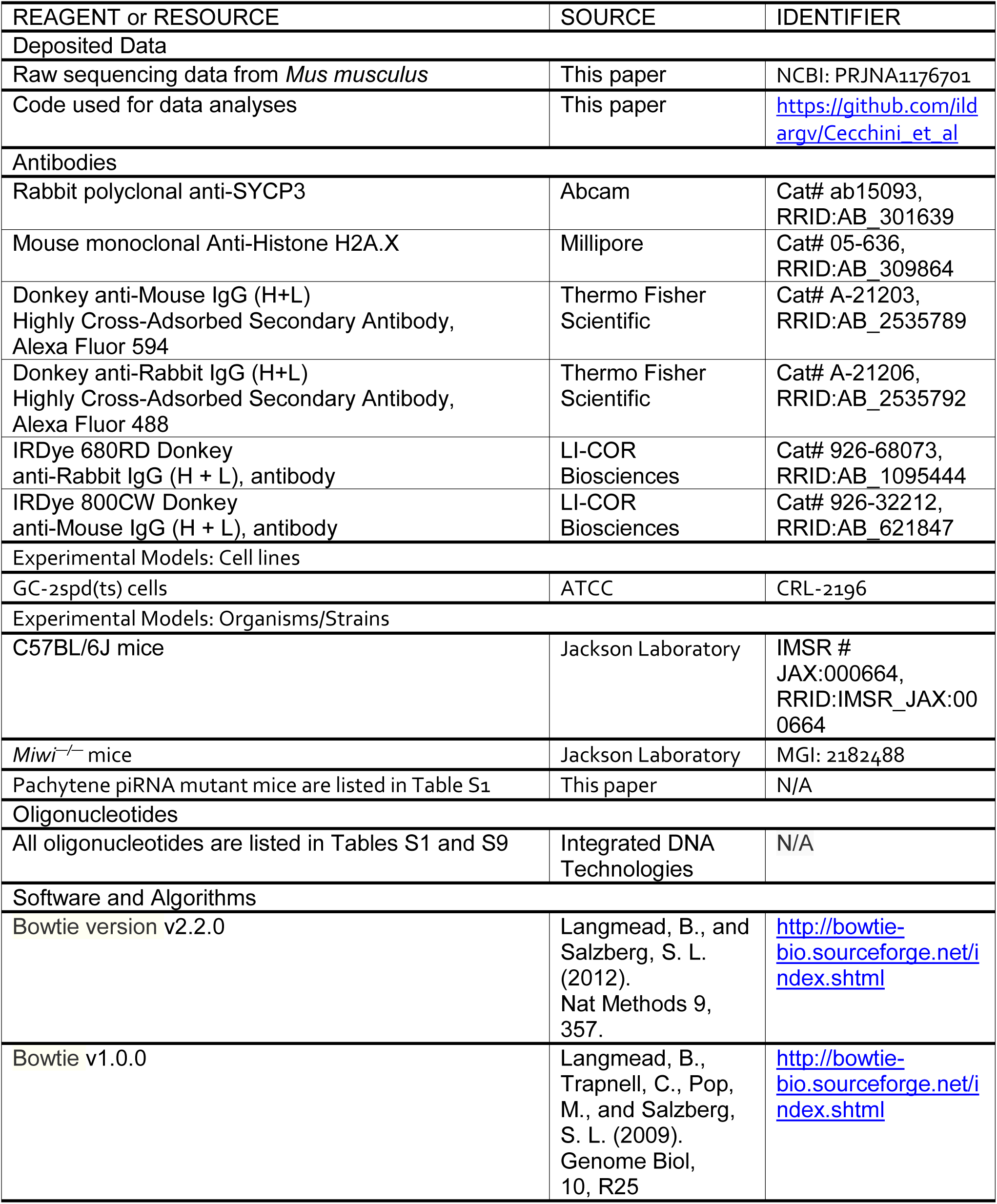
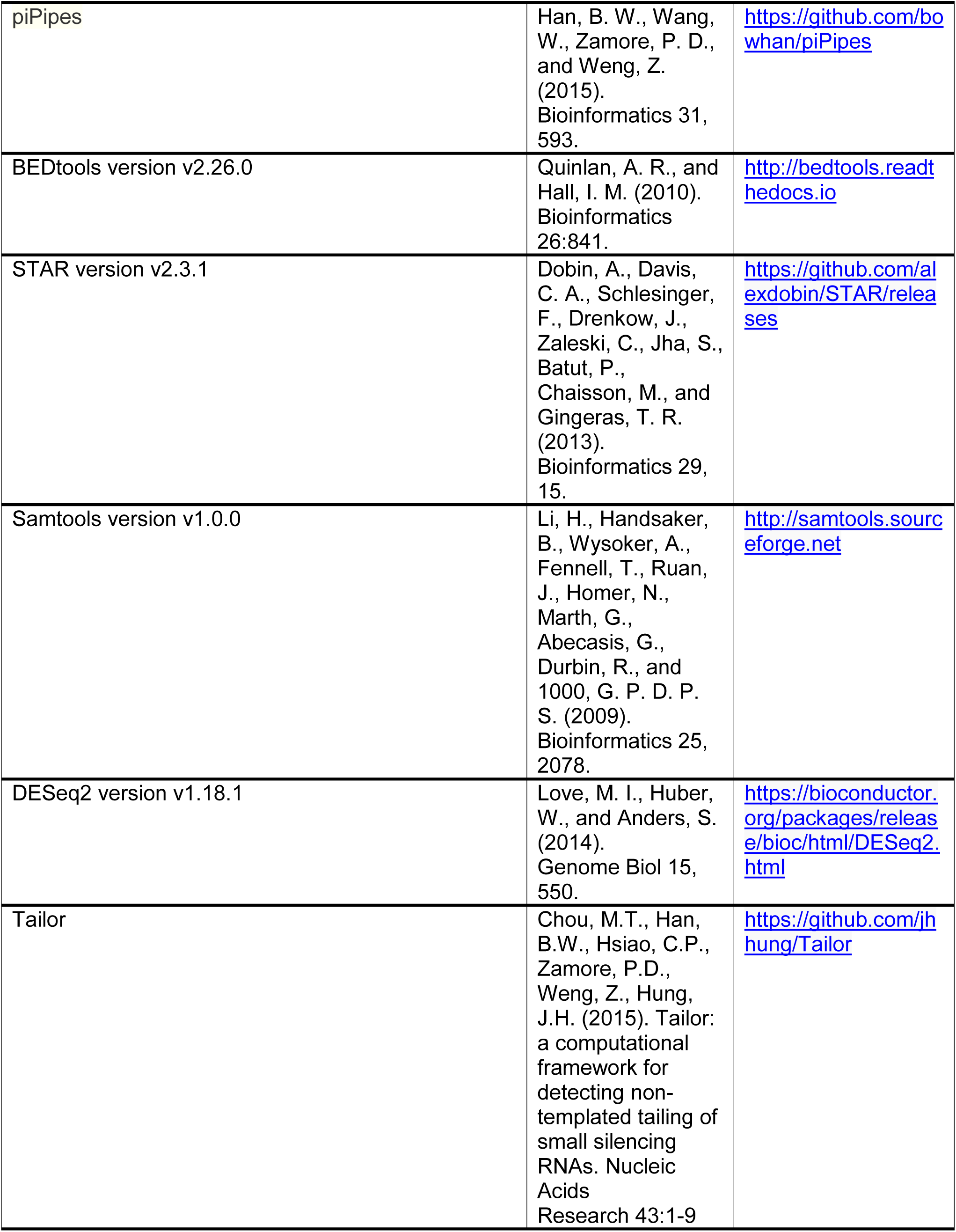
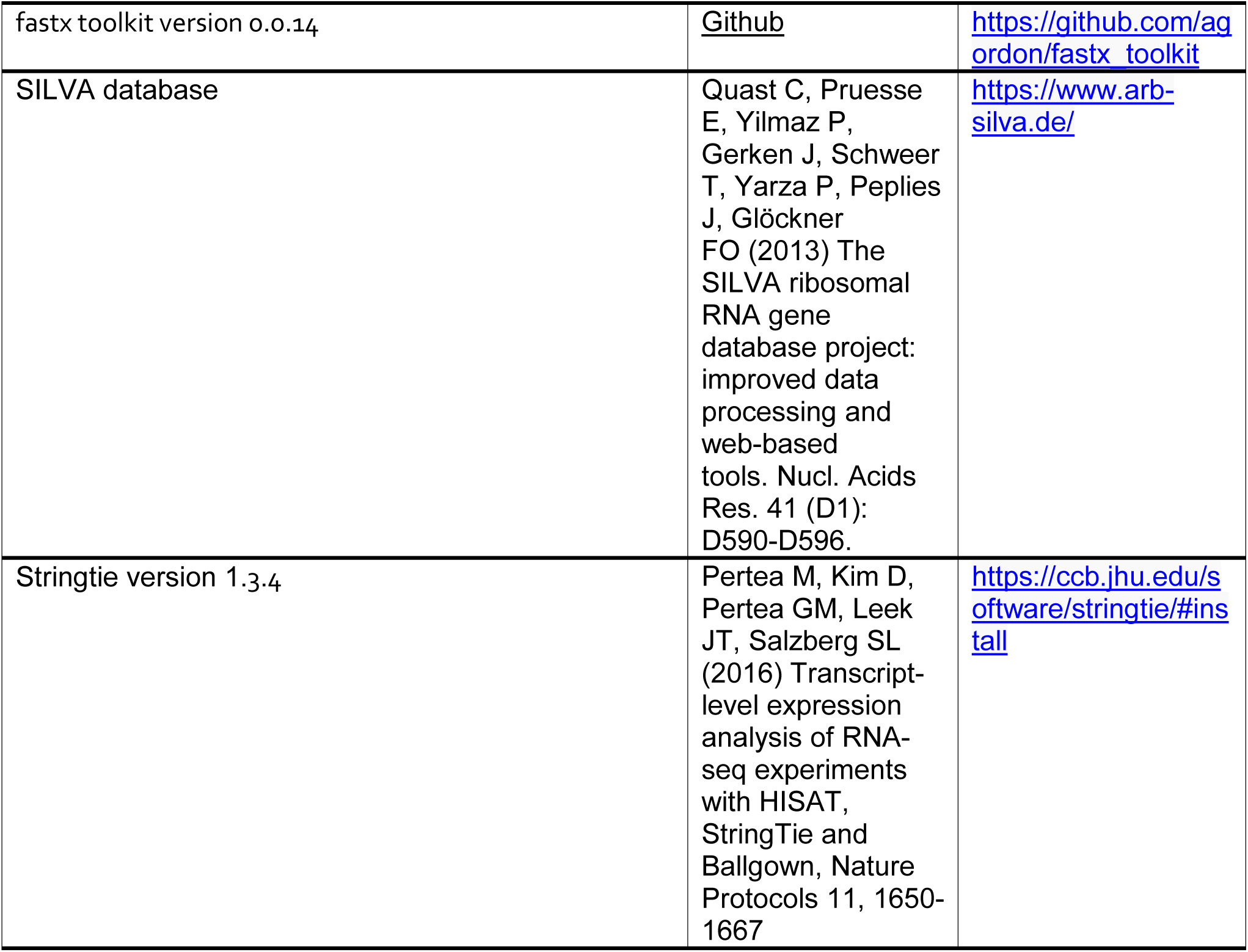
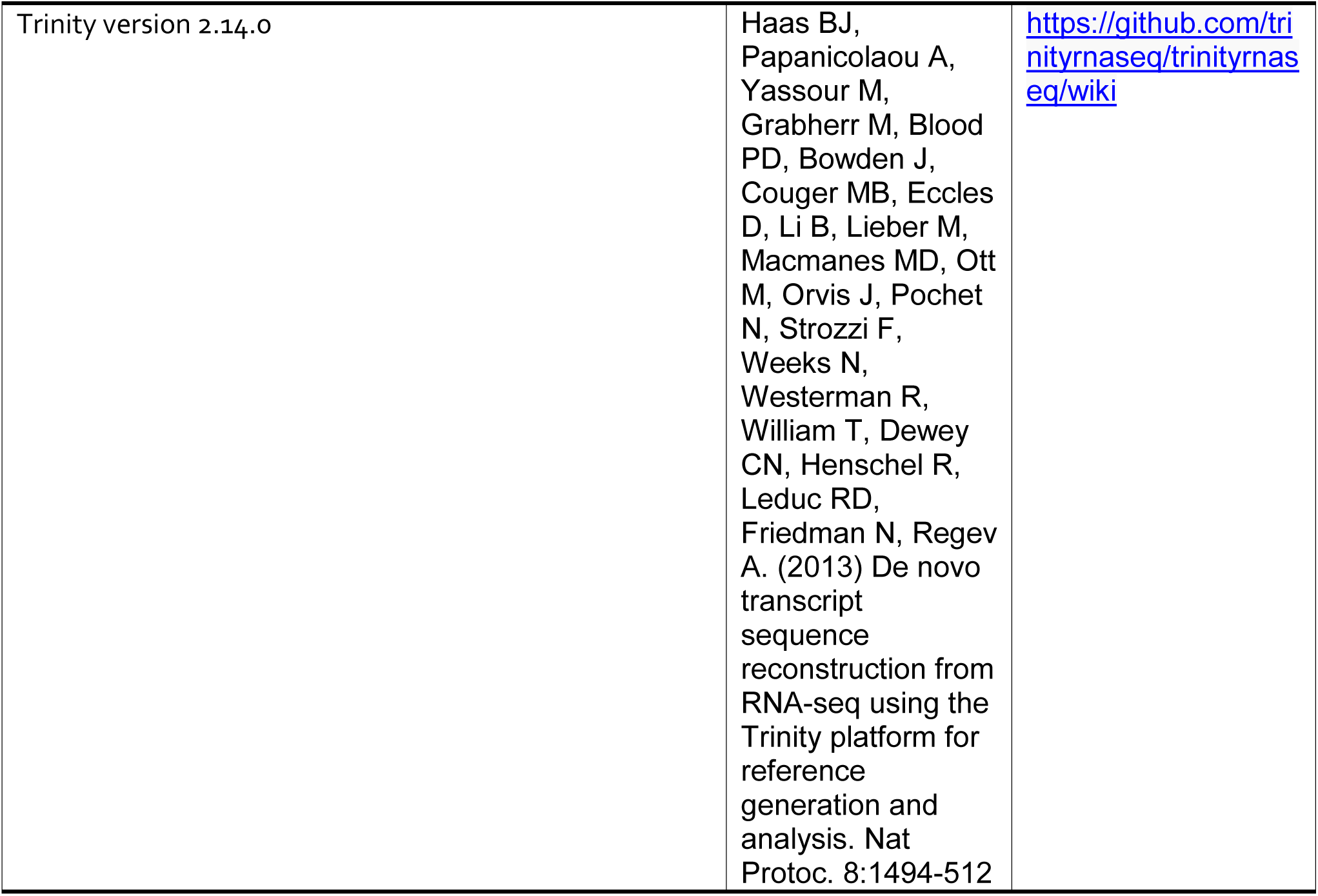

---

**Table S1.**
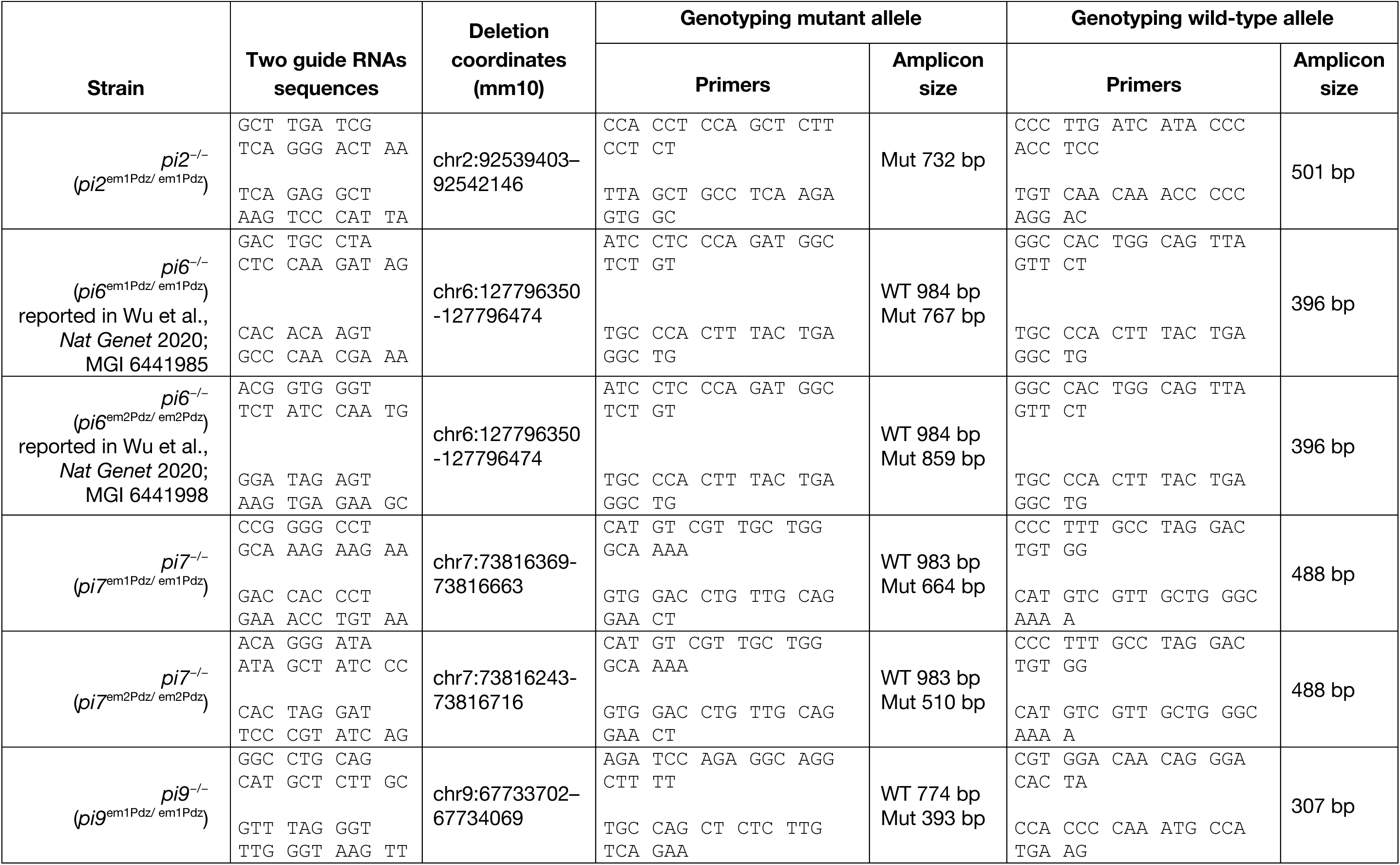
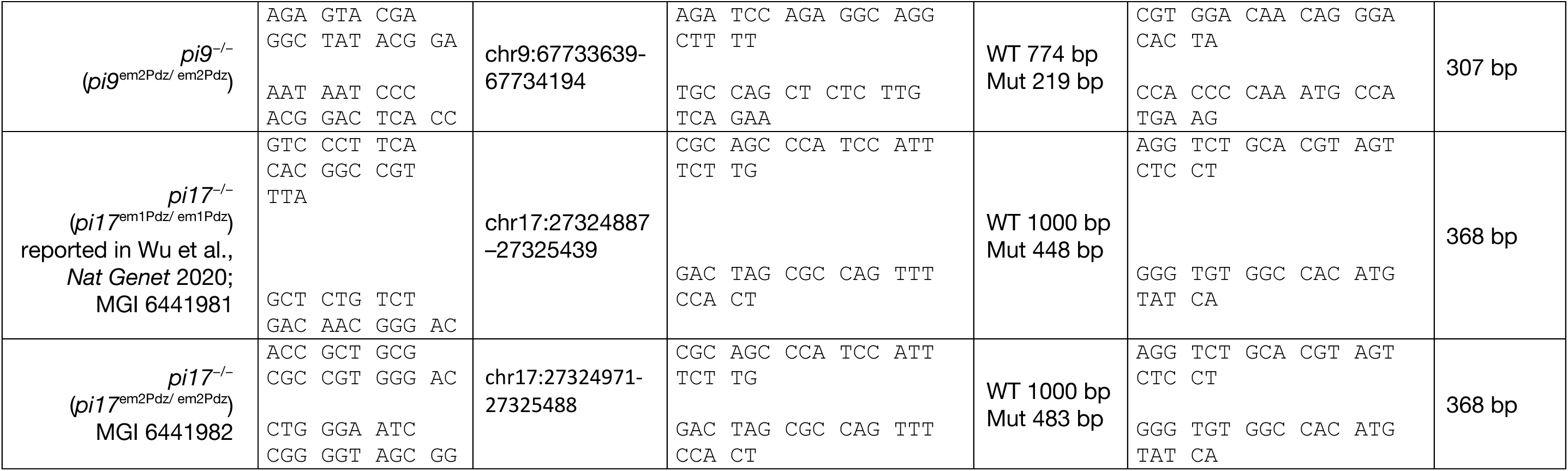
Mouse strains used in this study.

**Table S3A.**
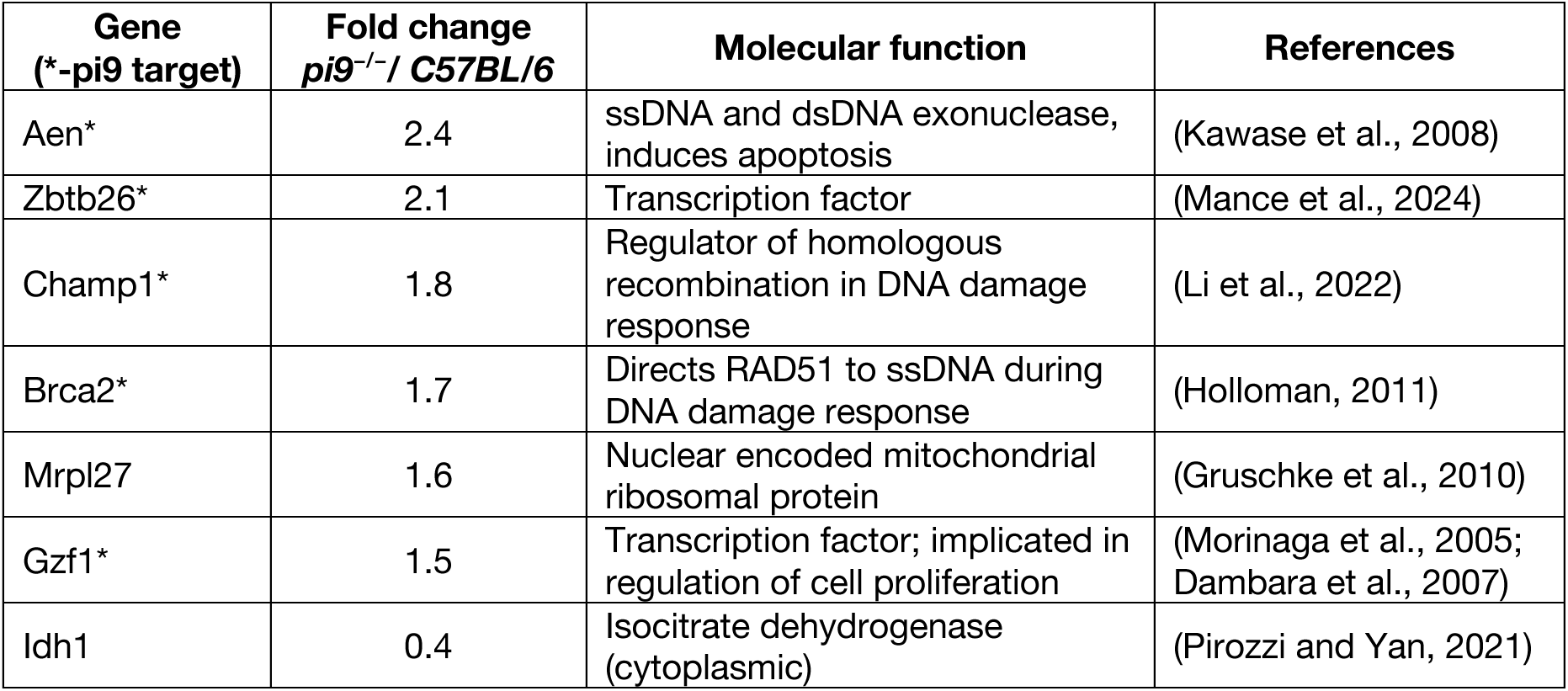
Molecular function of genes whose abundance changes significantly in *pi9^−/−^* primary spermatocytes (FDR<0.01).

**Table S3B.**
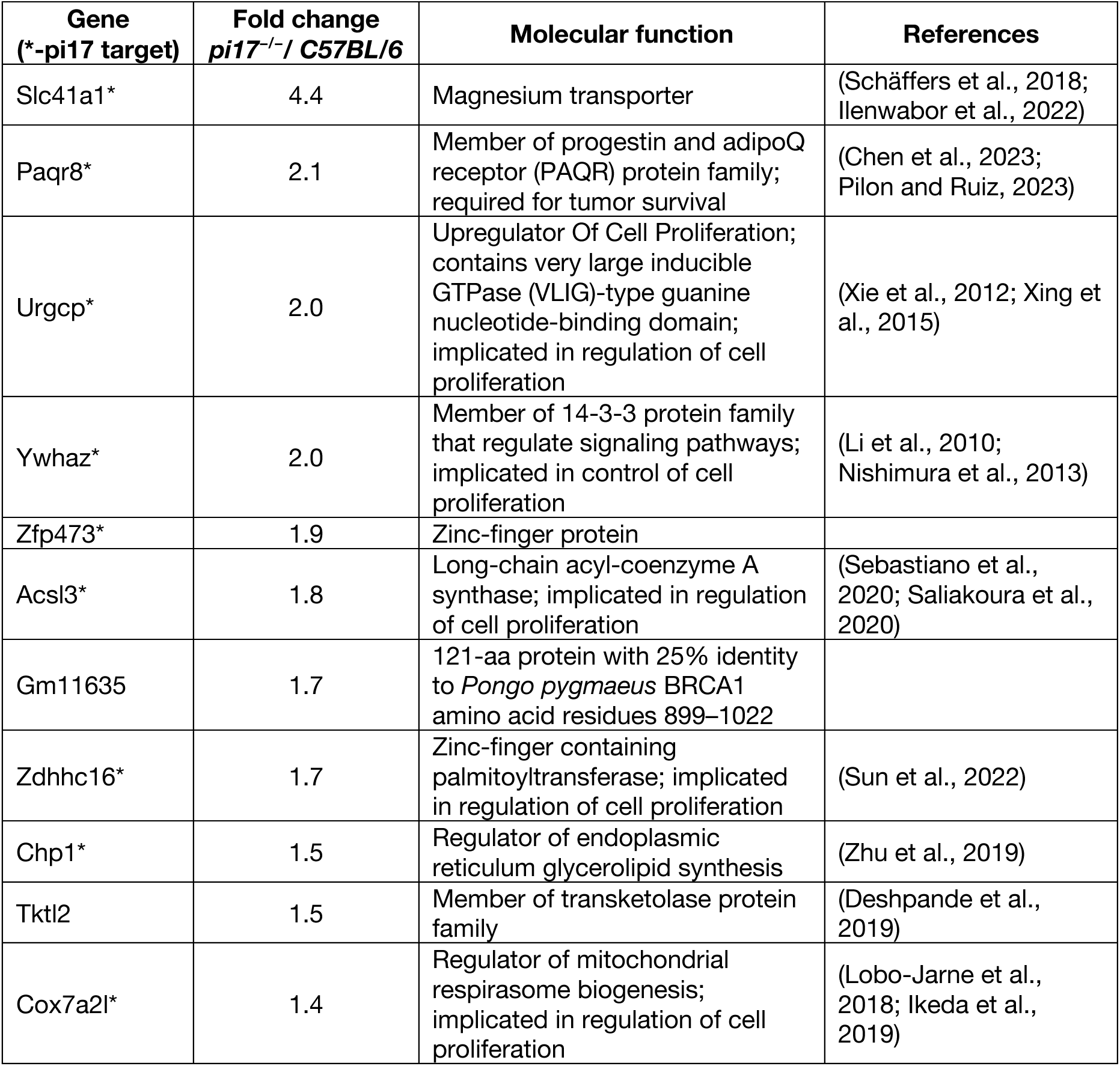
Molecular function of genes whose abundance changes significantly in *pi17^−/−^* primary spermatocytes (FDR<0.01).

**Table S8.**
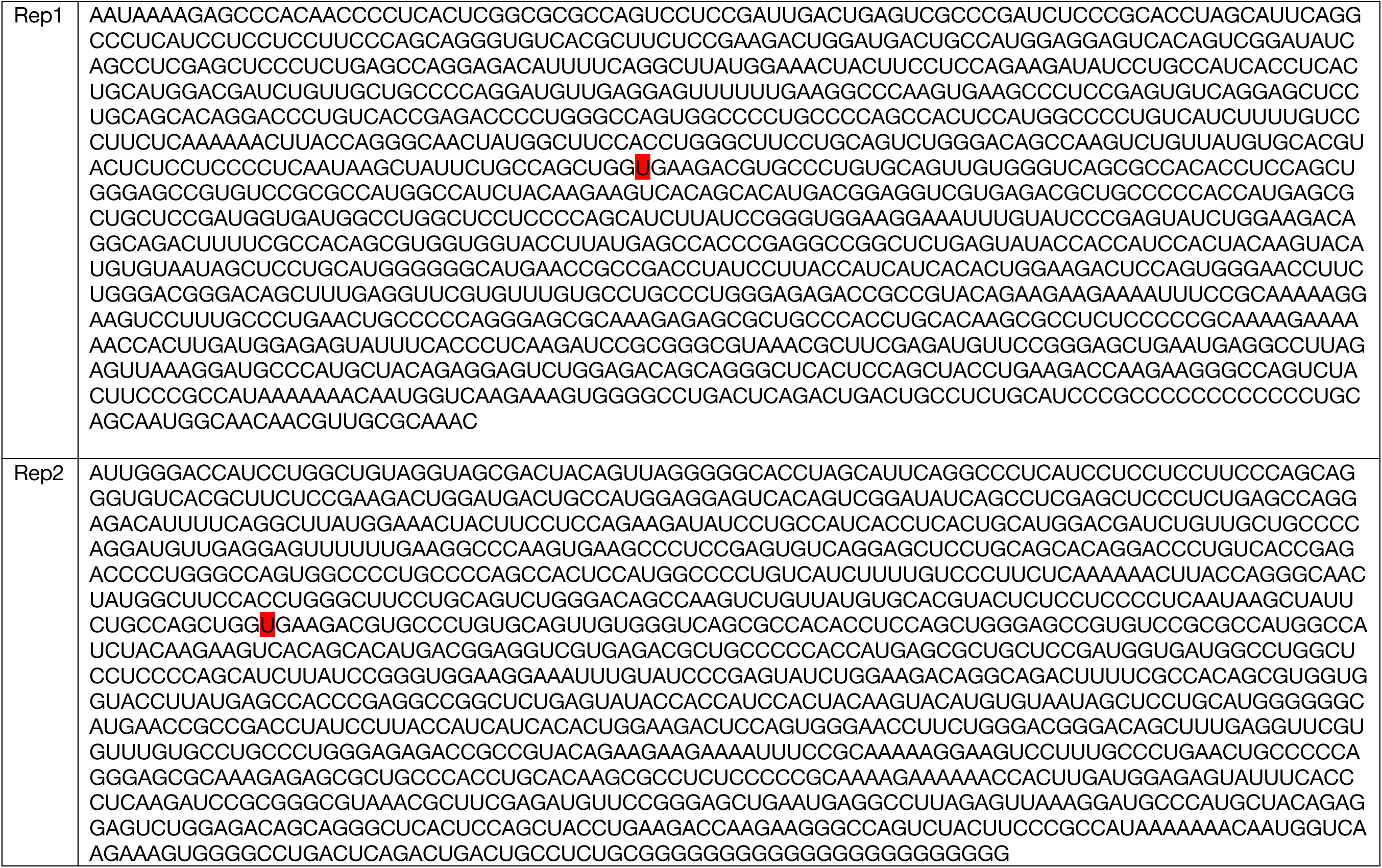
Sequence of *Trp53* mRNA assembled de novo with raw RNA-seq reads from GC-2spd(ts) cells using Trinity. Mutation that causes Ala-to-Val substitution is highlighted in red.

**Table S9.**
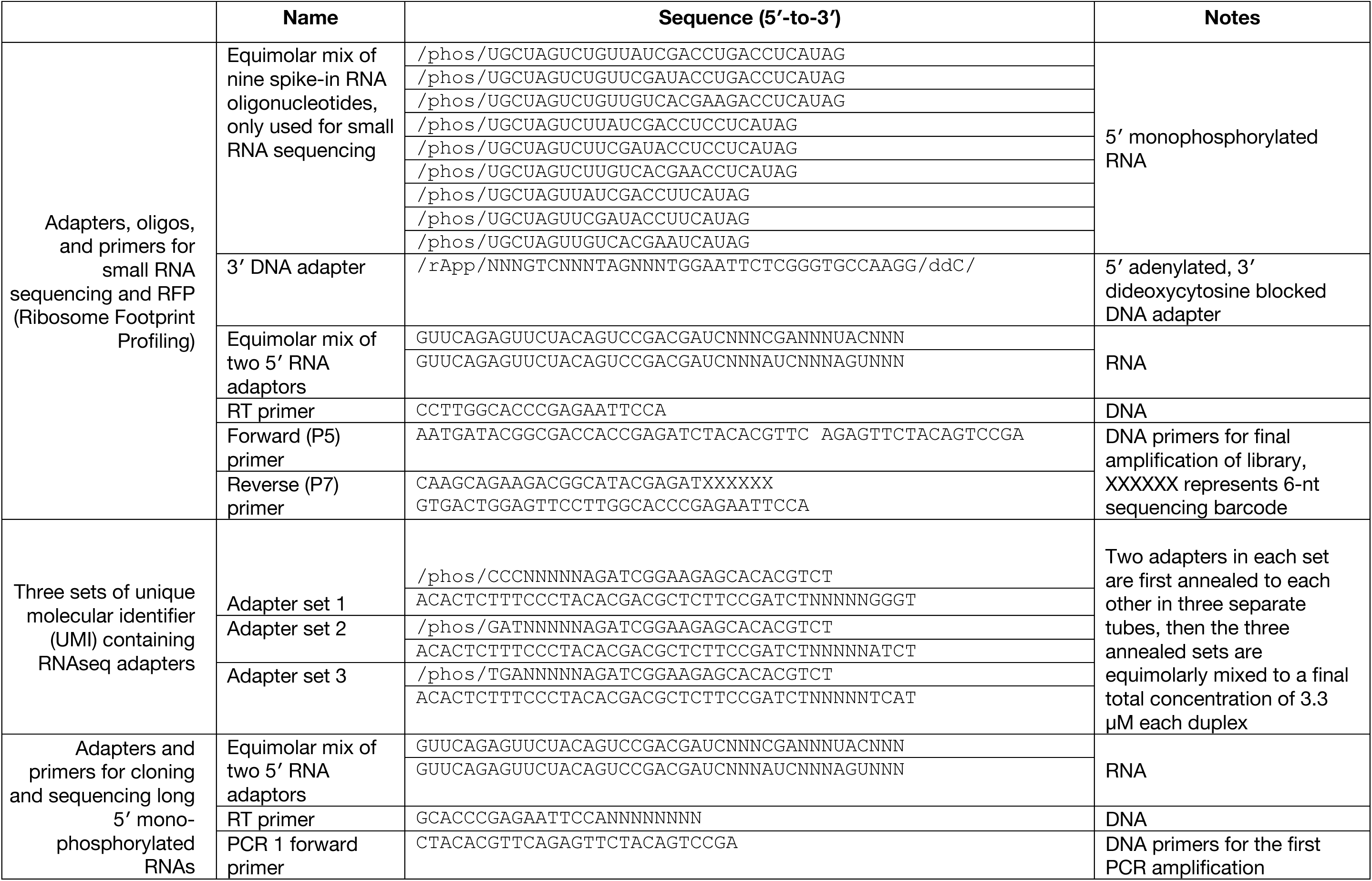
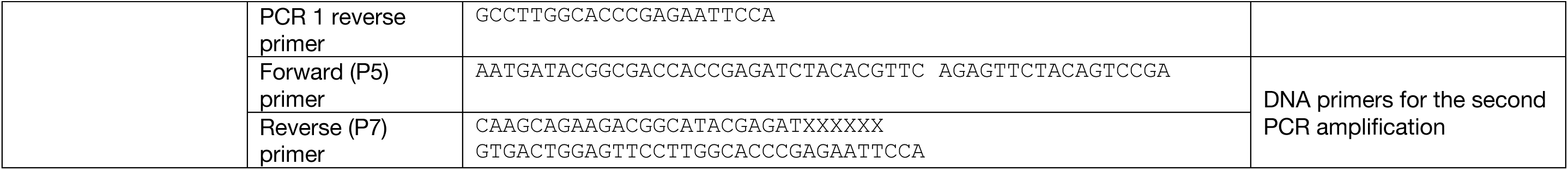
Sequences of oligonucleotides used in this study.

**Table S10.** Number of Primary Spermatocytes and GC-2spd(ts) cells and Amount of Spike-In Mix Used to Prepare Small RNA Sequencing Libraries.

| Genotype |  | Trial | Cell number | Amount of spike-in, attomol |
| --- | --- | --- | --- | --- |
| C57BL/6 data are from Gainetdinov et al, *Nature* 2023 | SRR21528503 | Rep1 | 31400 | 370 |
|  | SRR21528502 | Rep2 | 68900 | 4000 |
|  | SRR21528501 | Rep3 | 47200 | 3000 |
|  | SRR21528500 | Rep4 | 89100 | 4000 |
|  | SRR21528499 | Rep5 | 48900 | 3000 |
|  | SRR21528498 | Rep6 | 60300 | 4000 |
|  | SRR21528497 | Rep7 | 63100 | 4000 |
|  | SRR21528495 | Rep8 | 99700 | 4000 |
|  | SRR21528494 | Rep9 | 112500 | 4000 |
|  | SRR21528493 | Rep10 | 116400 | 4000 |
|  | SRR21528492 | Rep11 | 137000 | 4000 |
|  | SRR21528491 | Rep12 | 112000 | 4000 |
| *pi9<sup>-/-</sup>* |  | Rep1 | 41,600 | 3,000 |
|  |  | Rep2 | 47,100 | 3,000 |
|  |  | Rep3 | 45,600 | 3,000 |
|  |  | Rep4 | 59,600 | 4,000 |
|  |  | Rep5 | 71,200 | 4,000 |
|  |  | Rep6 | 95,800 | 4,000 |
|  |  | Rep7 | 56,000 | 4,000 |
| *pi17<sup>-/-</sup>* |  | Rep1 | 59,900 | 4,000 |
|  |  | Rep2 | 36,460 | 4,000 |
|  |  | Rep3 | 76,570 | 4,000 |
|  |  | Rep4 | 54,000 | 4,000 |
|  |  | Rep5 | 153,300 | 4,000 |
|  |  | Rep6 | 91,300 | 4,000 |
| *pi9<sup>-/-</sup>pi17<sup>-/-</sup>* |  | Rep1 | 40,000 | 4,000 |
|  |  | Rep2 | 36,000 | 4,000 |
|  |  | Rep3 | 33,500 | 4,000 |
| *pi6<sup>-/-</sup>* |  | Rep1 | 45,400 | 4,000 |
| GC-2spd(ts) |  | Rep1 | 100,000 | 400 |
|  |  | Rep2 | 100,000 | 400 |
|  |  | Rep3 | 100,000 | 400 |
|  |  | Rep4 | 100,000 | 400 |

